# Widely Used Variants of the Farquhar-von-Caemmerer-Berry Model Can Cause Errors in Parameter Estimation

**DOI:** 10.1101/2025.03.11.642611

**Authors:** Edward B. Lochocki, Justin M. McGrath

## Abstract

The Farquhar-von-Caemmerer-Berry (FvCB) model is the most widely-used mechanistic model of C_3_ net CO_2_ assimilation, and it plays a significant role in plant physiology, ecology, climate science, and Earth system modeling. As use of the model has grown, multiple variants have appeared across publications. Although many of these are commonly used, there has not been a detailed investigation of existing variants and their impacts on results and interpretations. Here we summarize the types of variants and their prevalence in the literature, and we present a comprehensive comparison of differences between them. A key finding is that a common variant that uses the minimum of assimilation rates rather than the minimum of carboxylation rates, which we call the “min-*A* variant,” makes different predictions than the original “min-*W* variant,” yet appears in approximately half of highly-cited publications and software tools that use the FvCB model. Another concern is that although leaf biochemistry restricts the range of CO_2_ partial pressures where limitations due to triose phosphate utilization (TPU) can occur, this restriction is commonly omitted from the model’s equations. Among other potential issues, these variations can introduce errors exceeding 20% when estimating photosynthetic parameter values from CO_2_ response curves. It is therefore important to be aware of this source of error when fitting the model, to avoid using the min-*A* variant, and to include the biochemically-derived CO_2_ threshold for TPU limitations.

**Central Theme of the Manuscript:** The Farquhar-von-Caemmerer-Berry model of CO_2_ assimilation plays a key role in plant research, but many publications use variants of the model that differ from the original and can potentially introduce errors in photosynthetic parameter estimates.

**Novel Results, Ideas, or Methods:** Using a literature survey, FvCB model variants are categorized, and some are found to make contradictory predictions. Comparisons against *A*-*C*_*i*_ curves show that the “min-*W* variant” exhibits the best performance, especially at low CO_2_ concentrations.

## 1. Introduction

The Farquhar-von-Caemmerer-Berry (FvCB) model for C_3_ net CO_2_ assimilation is a cornerstone of modern plant biology, ecology, and climate science, having been highly successful in explaining experimental measurements and making predictions at scales ranging from single cells to the entire globe (Farquhar et al., 2001; Von Caemmerer, 2013). Although commonly referred to by the names of the authors of a key 1980 publication (Farquhar et al., 1980), the FvCB model is nonetheless built on decades of research that predates 1980, and it has been improved by other researchers since its initial description (Yin et al., 2021). It is a mechanistic model based on a simplified version of the light-dependent reactions, the Calvin-Benson-Bassham (CBB) cycle, and the photorespiratory cycle, and it consists of a small set of equations for predicting net CO_2_ assimilation rates (Farquhar et al., 1980; Farquhar and von Caemmerer, 1982; Kirschbaum and Farquhar, 1984). In a plant physiology context, the many possible applications of the FvCB model include assessing the impact of elevated atmospheric CO_2_ concentrations on photosynthesis (Bernacchi et al., 2005), identifying limitations to photosynthetic CO_2_ assimilation (Busch and Sage, 2017), evaluating the genetic diversity of ribulose-1,5-bisphosphate carboxylase-oxygenase (Rubisco) carboxylation efficiency within a species (De Souza et al., 2020), and helping to guide strategies for developing sustainable and high-performing crops in the face of climate change (He and Matthews, 2023; Matthews et al., 2022; Wu et al., 2023; Yin and Struik, 2017).

The FvCB model works by separately calculating three potential rates of ribulose 1,5-bisphosphate (RuBP) carboxylation as catalyzed by Rubisco, which are limited by either Rubisco activity (*W*_*c*_), RuBP regeneration (*W*_*j*_), or triose phosphate utilization (TPU) (*W*_*p*_). The slowest potential rate is chosen as the actual RuBP carboxylation rate (*V*_*c*_). The net leaf-level CO_2_ assimilation rate (*A*_*n*_) can then be determined by subtracting CO_2_ losses due to RuBP oxygenation and non-photorespiratory CO_2_ release from the gains due to RuBP carboxylation. Mathematically, this process can be expressed by five equations:

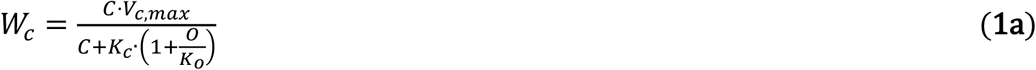

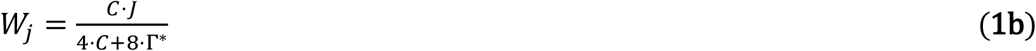

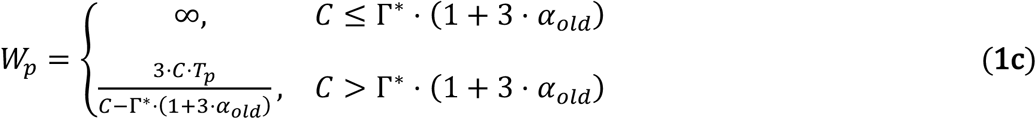

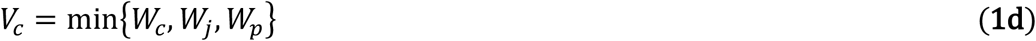

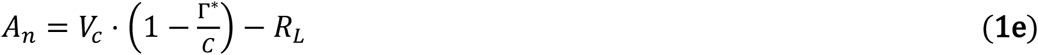

Here, *C* is the partial pressure of CO_2_ in the vicinity of Rubisco. Definitions for all the symbols used in these equations can be found in Table **1**. Although Equation 1 differs from early expressions of the FvCB model in its approach to TPU limitations (Farquhar and von Caemmerer, 1982; Kirschbaum and Farquhar, 1984), it follows the original reasoning for identifying the rate-limiting process, which we refer to as the “min-*W* approach.”

**Table 1:** Description of symbols used in this publication. Values marked with a superscript (a) are from the caption of Figure 2.6 in von Caemmerer (2000). The typical value for O in the table is the atmospheric value at sea level rounded to two significant figures. (b) By convention, the term “RuBP regeneration limitations” refers to the case where RuBP regeneration is limited primarily by electron transport, and possibly co-limited by the activity of CBB cycle enzymes such as sedoheptulose-1,7 bisphosphatase; limitations to RuBP regeneration imposed by the supply of inorganic phosphate or carbon are treated separately. (c) Consumption refers to the incorporation of inorganic phosphate into organic molecules. (d) Even when α_old_ is zero, one quarter of glycolate carbon is released as CO_2_ during glycine decarboxylation (Harley and Sharkey, 1991), so the overall fraction of glycolate carbon not returned to the chloroplast is (1 + 3 · α_old_)/4.

| Symbol | Meaning | Units | Range | Typical value |
| --- | --- | --- | --- | --- |
| $A_n$ | Net CO<sub>2</sub> assimilation rate | μmol m<sup>-2</sup> s<sup>-1</sup> | - | - |
| $A_c$ | Net CO<sub>2</sub> assimilation rate when RuBP carboxylation is limited by Rubisco activity | $\mu\text{mol m}^{-2} \text{s}^{-1}$ | - | - |
| $A_c\vert _{C=0}$ | The value of $A_c$ when $C = 0$ | $\mu\text{mol m}^{-2} \text{s}^{-1}$ | - | - |
| $A_d$ | Net CO<sub>2</sub> assimilation rate when RuBP carboxylation is limited by substantial Rubisco deactivation or RuBP depletion at low $C$ | $\mu\text{mol m}^{-2} \text{s}^{-1}$ | - | - |
| $A_j$ | Net CO<sub>2</sub> assimilation rate when RuBP carboxylation is limited by the rate of electron transport going to support RuBP regeneration<sup>(b)</sup> | $\mu\text{mol m}^{-2} \text{s}^{-1}$ | - | - |
| $A_j\vert _{C=0}$ | The value of $A_j$ when $C = 0$ | $\mu\text{mol m}^{-2} \text{s}^{-1}$ | - | - |
| $A_p$ | Net CO<sub>2</sub> assimilation rate when RuBP carboxylation is limited by the rate of inorganic phosphate release from TPU | $\mu\text{mol m}^{-2} \text{s}^{-1}$ | - | - |
| $C$ | Partial pressure of CO<sub>2</sub> in the vicinity of Rubisco | $\mu\text{bar}$ | $0 \leq C$ | - |
| $C_{cj}$ | The value of $C$ where $W_c = W_j$ | $\mu\text{bar}$ | $0 \leq C_{cj}$ | - |
| $C_c$ | Chloroplastic partial pressure of CO<sub>2</sub> | $\mu\text{bar}$ | $0 \leq C_c$ | - |
| $C_i$ | Intercellular partial pressure of CO<sub>2</sub> | $\mu\text{bar}$ | $0 \leq C_i$ | - |
| $C_i^*$ | The value of $C_i$ when $C_c = \Gamma^*$ | $\mu\text{bar}$ | $0 \leq C_i^*$ | - |
| $J$ | Potential rate of linear electron transport going to support RuBP regeneration at a given light intensity | $\mu\text{mol m}^{-2} \text{s}^{-1}$ | $0 \leq J$ | 170<sup>(a)</sup> |
| $J_{eq}$ | The value of $J$ where $A_c\vert _{C=0} = A_j\vert _{C=0}$ | $\mu\text{mol m}^{-2} \text{s}^{-1}$ | $0 \leq J_{eq}$ | - |
| $J_d$ | The value of $J$ where the denominator of Supplemental Equation A5 is zero | $\mu\text{mol m}^{-2} \text{s}^{-1}$ | $0 \leq J_d$ | - |
| $J_n$ | The value of $J$ where the numerator of Supplemental Equation A5 is zero | $\mu\text{mol m}^{-2} \text{s}^{-1}$ | $0 \leq J_n$ | - |
| $K_c$ | Michaelis-Menten constant for CO<sub>2</sub> | $\mu\text{bar}$ | $0 < K_c$ | 259<sup>(a)</sup> |
| $K_o$ | Michaelis-Menten constant for O<sub>2</sub> | mbar | $0 < K_o$ | 179<sup>(a)</sup> |
| $O$ | Partial pressure of O<sub>2</sub> in the vicinity of Rubisco | mbar | $0 \leq O$ | 210 |
| $Q_{in}$ | Incident photosynthetically active flux density | $\mu\text{mol m}^{-2} \text{s}^{-1}$ | $0 \leq Q_{in}$ | - |
| $R_L$ | Rate of non-photorespiratory CO<sub>2</sub> release in the light | $\mu\text{mol m}^{-2} \text{s}^{-1}$ | $0 \leq R_L$ | 1<sup>(a)</sup> |
| $R_{pc}$ | Net rate of inorganic phosphate consumption in the chloroplast due to photosynthesis and photorespiration<sup>(c)</sup> | $\mu\text{mol m}^{-2} \text{s}^{-1}$ | - | - |
| $T_p$ | Potential rate of TPU | $\mu\text{mol m}^{-2} \text{s}^{-1}$ | $0 < T_p$ | 11.8<sup>(a)</sup> |
| $V_c$ | RuBP carboxylation rate | $\mu\text{mol m}^{-2} \text{s}^{-1}$ | $0 \leq V_c$ | - |
| $V_{c,max}$ | Maximum rate of Rubisco carboxylation activity | $\mu\text{mol m}^{-2} \text{s}^{-1}$ | $0 \leq V_{c,max}$ | 100<sup>(a)</sup> |
| $V_o$ | RuBP oxygenation rate | $\mu\text{mol m}^{-2} \text{s}^{-1}$ | $0 \leq V_o$ | - |
| $W_c$ | RuBP carboxylation rate limited by Rubisco activity | $\mu\text{mol m}^{-2} \text{s}^{-1}$ | $0 \leq W_c$ | - |
| $W_d$ | RuBP carboxylation rate limited by substantial Rubisco deactivation or RuBP depletion at low $C$ | $\mu\text{mol m}^{-2} \text{s}^{-1}$ | $0 \leq W_d$ | 0 |
| $W_j$ | RuBP carboxylation rate limited by the rate of electron transport going to support RuBP regeneration<sup>(b)</sup> | $\mu\text{mol m}^{-2} \text{s}^{-1}$ | $0 \leq W_j$ | - |
| $W_p$ | RuBP carboxylation rate limited by the rate of inorganic phosphate release from TPU | $\mu\text{mol m}^{-2} \text{s}^{-1}$ | $0 \leq W_p$ | - |
| $\alpha_{old}$ | Fraction of remaining glycolate carbon not returned to the chloroplast after accounting for carbon released as $\text{CO}_2$<sup>(d)</sup> | dimensionless | $0 \leq \alpha_{old} \leq 1$ | 0<sup>(a)</sup> |
| $\alpha_G$ | Fraction of glycolate carbon exported from the photorespiratory pathway as glycine | dimensionless | $0 \leq \alpha_G \leq 1$ | 0 |
| $\alpha_S$ | Fraction of glycolate carbon exported from the photorespiratory pathway as serine | dimensionless | $0 \leq \alpha_S \leq \frac{3}{4}$ | 0 |
| $\alpha_T$ | Fraction of glycolate carbon exported from the photorespiratory pathway as $\text{CH}_2\text{-THF}$ | dimensionless | $0 \leq \alpha_T \leq \frac{1}{2}$ | 0 |
| $\Gamma^*$ | $\text{CO}_2$ compensation point in the absence of non-photorespiratory $\text{CO}_2$ release | $\mu\text{bar}$ | $0 < \Gamma^*$ | 38.6<sup>(a)</sup> |
| $\Delta_c$ | Net rate of carbon production during the reduction of PGA to unbound RuBP | $\mu\text{mol m}^{-2} \text{s}^{-1}$ | - | - |

The conditional form of Equation **1c** ensures that *W*_*p*_ ≥ 0 (as is necessary for a carboxylation rate) and that TPU cannot impose a limit on *V*_*c*_ when *C* ≤ Γ^*^ · (1 + 3 · *α*_*old*_). This lower *C* threshold for TPU limitations will be discussed later in more detail. Note that a more complex version of the model separately includes glycolate carbon leaving the photorespiratory pathway as glycine, serine, or 5,10-methylene-tetrahydrofolate (CH_2_-THF), requiring three *α* parameters instead of one: *α*_*G*_, *α*_*S*_, and *α*_*T*_ (Busch, 2020; Busch et al., 2018). Using the *α*_*old*_ model provides simplicity as it has fewer parameters, and thus it can more easily be applied. However, the biochemical basis for the three *α* version is better established, and it would be appropriate when the detail is necessary and the additional parameter values can be estimated. Here, we use the simpler *α*_*old*_ version for simplicity when comparing the other variants because the Busch *et al*. updates do not affect the general conclusions presented here (Supplemental Section **S1.2**).

Most applications of the FvCB model in scientific research require knowledge of the values of key parameters such as *V*_*c,max*_, *J*, and *T*_*p*_; for example, parameter values may be used to characterize groups of plants or as inputs to computational models. These parameter values can be estimated from experimentally measured CO_2_ response curves, which are obtained by using a gas exchange system to record *A*_*n*_ and *C*_*i*_ as the CO_2_ concentration around a leaf is varied under otherwise constant conditions of incident light, humidity, and temperature. By comparing the measured curve against predictions from the FvCB model made using *C* = *C*_*i*_ or, if possible, *C* = *C*_*c*_, it is possible to find values of the model parameters that best reproduce the measured assimilation rates. Many approaches to curve fitting can be found in the literature, some of which include additional types of data that can be used to estimate *C*_*c*_, such as chlorophyll fluorescence (Bellasio et al., 2016; Duursma, 2015; Gu et al., 2010; Lochocki et al., 2025, 2025; Moualeu-Ngangue et al., 2017; Sharkey et al., 2007; Stinziano et al., 2021; Wang et al., 2017; Xiao et al., 2021).

As usage of the model for curve fitting and other purposes has increased, numerous additions and changes have occurred across scientific publications and software tools. Although many researchers are familiar with the uses and drawbacks of these variants, a detailed comparison would be valuable for those learning about the model, with the aim of providing insight into which variants may be appropriate for particular scenarios. Here we summarize FvCB model variants and key differences between them (Table **2** and Supplemental Section **S1**). In some cases, the choice of a variant may depend on the situation, but we also demonstrate that some variants provide no benefits and can produce results that contradict measurements. One such group of variants replaces Equations **1d** and **1e** by a single equation calculating *A*_*n*_ as the minimum of three potential net assimilation rates (Figure **1a**). As described in detail below, this approach, which we refer to as the “min-*A* approach,” alters the model’s predictions for *A*_*n*_ when *C* < Γ^*^ such that they disagree with measurements. Another issue is that although TPU limitations are understood to only occur at high *C* (Harley and Sharkey, 1991; Sharkey, 2019; Sharkey et al., 2007), the explicit lower *C* threshold for TPU included in Equation **1c** is often omitted, leading to situations where the model incorrectly predicts TPU to be limiting. The prevalence of these model variations in the literature and the potential consequences of using them in parameter estimates have not been explored.

**Table 2:** Classification scheme for FvCB model equations or computer code. To fully describe a variant, its approach in each category must be specified; for example, 0-α+NT+min-A+NFL is a variant with no α parameters or TPU limitations where the limiting process is chosen using the min-A approach without any other constraints. Note that not all combinations are possible, and that some are special cases of others; for example, 0-α and 1-α are equivalent when α_old_ is zero.

| Category | Approach | Abbreviation |
| --- | --- | --- |
| Treatment of carbon leaving the photorespiratory pathway | Not considered | 0-α |
| | $\alpha_{old}$ | 1-α |
| | $\alpha_G$ and $\alpha_S$ | 2-α |
| | $\alpha_G$, $\alpha_S$, and $\alpha_T$ | 3-α |
| CO<sub>2</sub> threshold for TPU limitations | No TPU limitations | NT |
|  | No explicit TPU threshold | NTT |
|  | Fixed TPU threshold | FTT |
|  | Biochemically-derived TPU threshold | BTT |
| Choice of limiting process | Minimum potential carboxylation rate | min-$W$ |
| | Minimum potential net CO<sub>2</sub> assimilation rate | min-$A$ |
| Additional constraints on choice of limiting process | No forced limitations | NFL |
|  | Rubisco limitations forced at low CO<sub>2</sub> | FRL |

**Figure 1:**
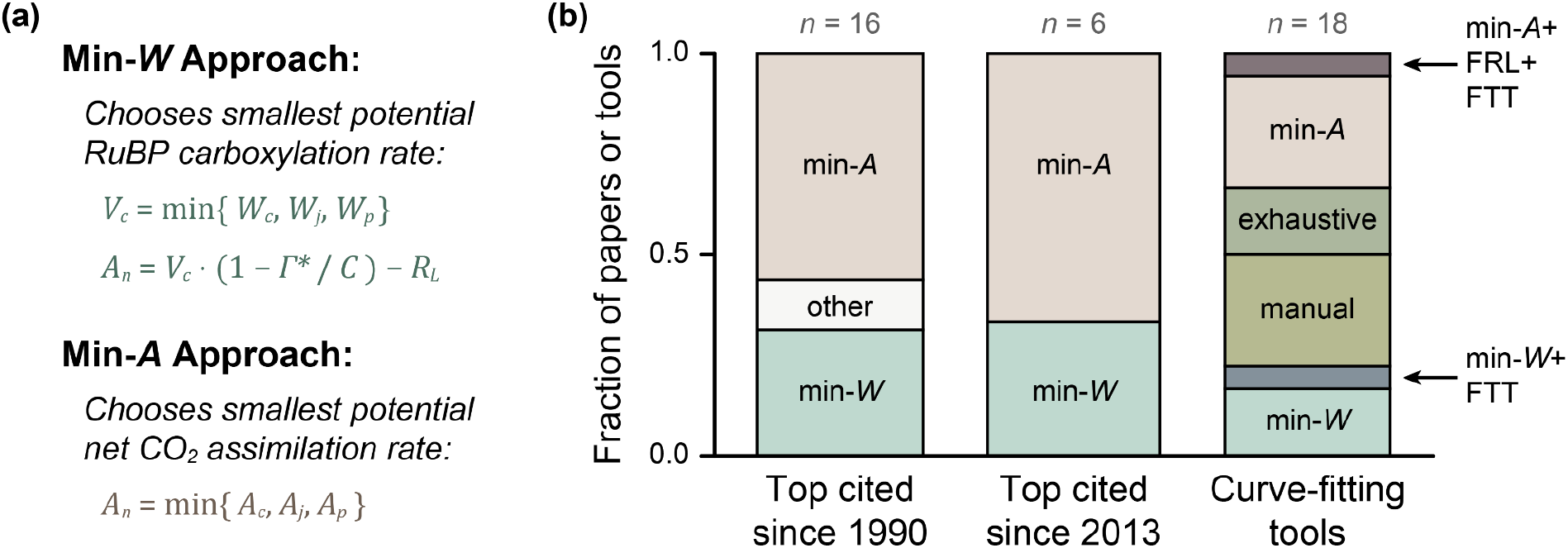
Description of the min-W and min-A approaches, and an estimate of their relative prevalence in the scientific literature. (**a**) Overview of the essential difference between the min-W and min-A approaches. The carboxylation rates W_c_, W_j_, and W_p_ that appear in the min-W approach can be calculated using Equation **1**, while the assimilation rates A_c_, A_j_, and A_p_ that appear in the min-A approach can be calculated using Equation **2**. (**b**) Fraction of highly cited papers (or tools for estimating FvCB model parameter values from CO_2_ response curves) that use the min-W approach, the min-A approach, or other approaches (Supplemental Section **S2**).

While it is difficult or impossible to determine how often researchers use a particular variant, estimates can be made by surveying peer-reviewed publications (Supplemental Section **S2**). The min-*A* approach first appears in the literature around 1990, with Collatz et al. (1990) and Collatz et al. (1991) being the earliest examples found in the survey. Out of the highest-cited papers published since 1990 that discuss the FvCB model equations, slightly more than half use the min-*A* approach (Figure **1b**). The min-*A* approach was identified as an “incorrect form” of the FvCB model in a 2010 publication (Gu et al., 2010), but its problems were not explained or demonstrated, and thus highly-cited publications have continued to use it with roughly the same frequency since 2013 (Figure **1b**). Likewise, the min-*A* approach is more likely to be used in popular software tools for estimating FvCB model parameters from CO_2_ response curves (Figure **1b**). Among the publications identified in this survey, only two include an explicit lower *C* threshold for TPU limitations: Gu et al. (2010) and Lochocki et al. (2025).

The survey of software tools also reveals the presence of two additional variations, which we refer to as the “forced Rubisco limitation” (FRL) and “fixed TPU threshold” (FTT) modifications (Table **2**). The FRL modification enforces Rubisco limitations at low *C* such that *A*_*n*_ = *A*_*c*_ for *C* ≤ Γ^*^. The FTT modification prevents TPU limitations for *C* below a fixed threshold value, rather than the one in Equation **1c**, which is determined from leaf biochemistry. These modifications address some, but not all, biological and numerical issues related to other variants of the FvCB model equations, as described below.

Although the literature survey presented here is limited (and does not include any software tools used in climate or agricultural modeling), it strongly suggests that many researchers working with biochemical models of C_3_ net CO_2_ assimilation have seen or used the min-*A* approach rather than the original min-*W* approach. It may even be possible that the min-*A* approach is more prevalent overall. Many researchers may also be unaware of the lower *C* threshold for TPU limitations. Given the central role of the FvCB model in several fields, it is necessary to elucidate the differences between the min-*W* and min-*A* approaches (with or without the FRL and FTT modifications) and to quantify the errors resulting from use of the min-*A* approach to estimate photosynthetic parameter values.

It might be argued that the differences between the min-*W* and min-*A* approaches are immaterial because all variants are incorrect when *C* < Γ^*^ due to Rubisco deactivation and RuBP depletion, processes that are not explicitly included in the FvCB model. Based on this reasoning, some researchers may intentionally avoid making gas exchange measurements or performing curve fits at low *C*. Yet, the ability to fit CO_2_ response curves for *C* < Γ^*^ has not been systematically tested. Thus, there is an opportunity to better understand net CO_2_ assimilation at low *C* by first expanding the FvCB model to include Rubisco deactivation and RuBP depletion and then comparing the min-*W* and min-*A* approaches against measured assimilation rates for *C* < Γ^*^.

The objectives of this study are to (1) describe the key differences between FvCB model variants, (2) establish the biochemical basis of the lower *C* threshold for TPU, (3) develop a simple way to include Rubisco deactivation and RuBP depletion in the FvCB framework, and (4) test the validity of the min-*W* and min-*A* approaches by fitting CO_2_ response curves that extend to *C* < Γ^*^. Our findings indicate that the min-*A* approach produces predictions that contradict observations, that the unrealistic predictions of the min-*A* approach cannot be fixed through the FRL and FTT modifications, and that the min-*A* approach cannot easily be extended to include Rubisco deactivation or RuBP depletion. In particular, the curve fits show that the min-*W* approach is able to represent C_3_ net CO_2_ assimilation at low *C* while the min-*A* approach cannot, leading to underestimates of *V*_*cmax*_ and *J* by up to 23% and 12%, respectively. Although errors are unlikely to be as severe in most situations, the min-*A* approach did not provide better predictions in any situation.

## 2. Description

### 2.1. The Min-W Variant

We categorize variations in FvCB model equations along several dimensions, such as the method used to determine the rate-limiting process, or the treatment of glycolate carbon leaving the photorespiratory pathway (Table **2**). Each dimension has several possible approaches, and each combination of possible approaches defines a variant. Equation **1** uses one *α* parameter (1-*α*), chooses a minimal carboxylation rate (min-*W*), uses the biochemically derived threshold for TPU limitations (BTT), and does not force Rubisco limitations at low *C* (NFL). Thus, it would be described as the 1-*α*+BTT+min-*W*+NFL variant. We refer to it as “the min-W variant” for brevity, since it is a representative example of the broader group of variants that use the min-*W* approach.

Here we note that within the time scale of a typical gas exchange measurement, the main environmental influences on the potential carboxylation rates are *C, Q*_*in*_, *O*, and temperature.

Note that some of this influence is not immediately obvious; for example, Γ^*^ depends on *O*, and *J* depends on *Q*_*in*_ and other factors (von Caemmerer, 2000). For simplicity, we assume constant temperature and treat Γ^*^ and *J* as inputs to the model, especially since several different approaches are available for calculating *J* from *Q*_*in*_ (Walker et al., 2021). As *C, J*, and *O* vary, the slowest carboxylation rate may change. Changes in the rate-limiting process as *C* increases from zero will be discussed later in more detail (Section **3.2**).

### 2.2. The Min-A Variant

When RuBP carboxylation is Rubisco-limited, the corresponding net assimilation rate (called *A*_*c*_ rather than *A*_*n*_ to indicate the rate-limiting process) can be found by setting *V*_*c*_ = *W*_*c*_ in Equation **1e** and replacing it with the expression in Equation **1a**:

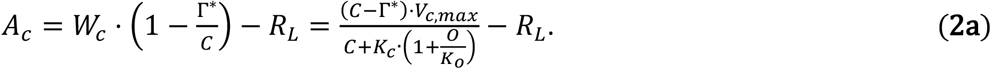

Likewise, when RuBP carboxylation is limited by RuBP regeneration or TPU, the corresponding net assimilation rates are given by analogous equations:

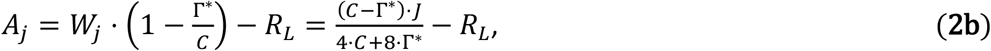

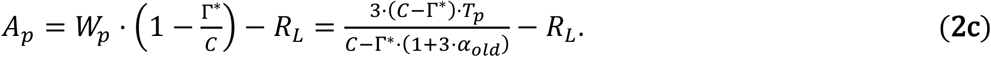

In the min-*A* approach, *A*_*c*_, *A*_*j*_, and *A*_*p*_ are considered to be potential net CO_2_ assimilation rates and the actual net CO_2_ assimilation rate is calculated as the smallest of the three, in analogy with Equations **1d** and **1e** from the min-*W* variant:

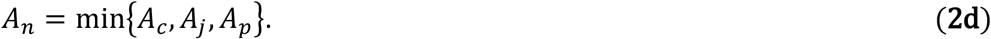

Specifically, Equation **2** describes the 1-*α*+NTT+min-*A*+NFL variant; we refer to this as the “min-*A* variant” for brevity, since it is a representative example of the broader group of variants that use the min-*A* approach.

Thus, the essential difference between Equations **1** and **2** is that the min-*W* variant chooses a minimal potential carboxylation rate, while the min-*A* variant chooses a minimal potential net CO_2_ assimilation rate (Figure **1a**). Another difference is that Equation **1c** restricts TPU limitations to *C* > Γ^*^ · (1 + 3 · *α*_*old*_), while Equation **2c** makes no such restriction. Many sources presenting the min-*W* approach also omit this restriction. Nevertheless, unreasonable results such as negative carboxylation rates and divergent behavior can occur in either variant when this restriction is not included; these issues have been noticed before and the FTT modification has been used to address them, although some issues remain when using a fixed TPU threshold (Section **3.3**).

## 3. Results

### 3.1 Minimal Carboxylation Rates Are Not Equivalent to Minimal Assimilation Rates

There are two irreconcilable differences between the min-*W* and min-*A* variants (Figure **2**).

**Figure 2:**
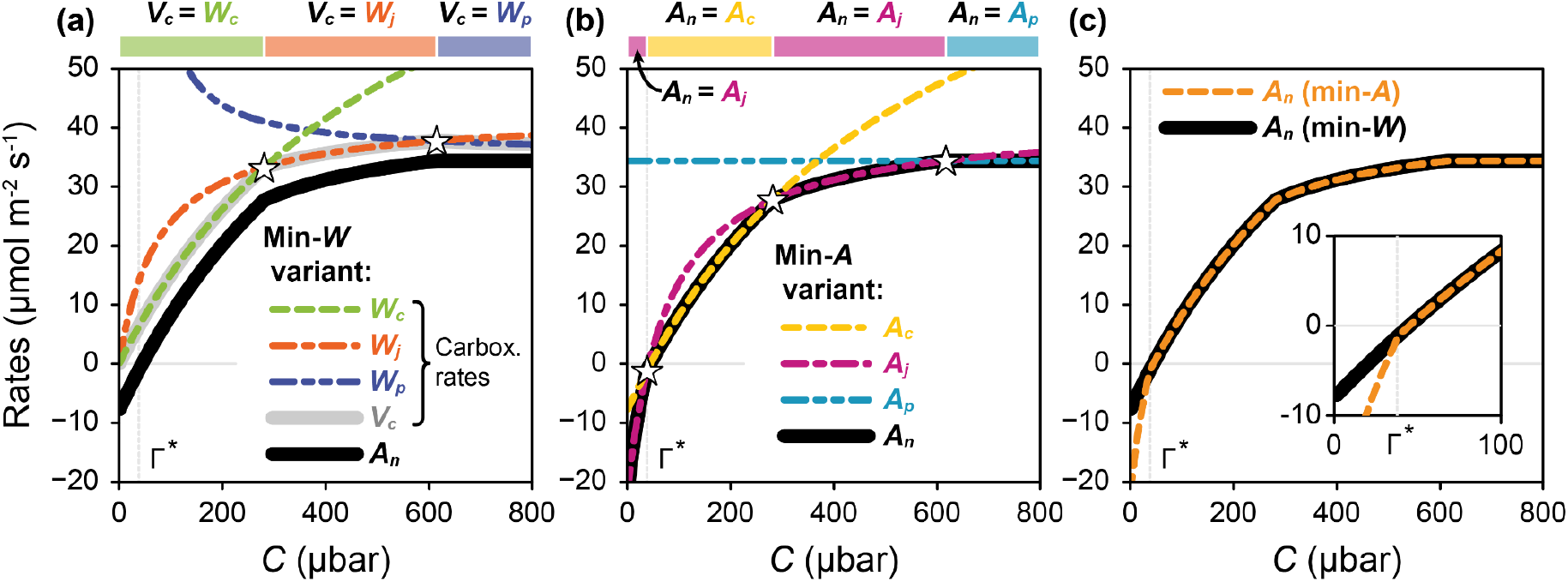
Comparison of outputs from the min-W and min-A variants for identical input parameter values with α_old_ = 0. (**a**,**b**) A_n_ (thick solid lines) as determined from potential carboxylation or assimilation rates (dashed lines) using Equations **1** and **2**, respectively. Crossover points where the rate-limiting process changes are marked with white stars. Solid rectangles indicate ranges of C where carboxylation or assimilation is limited by Rubisco, RuBP regeneration, or TPU. (**c**) Direct comparison of the variants across a range of C values. Inset: close-up showing the difference between the variants when C < Γ^*^. All parameters were set to the values specified in Table **1**.

1. When *C* < Γ^*^, the two variants always disagree about the rate-limiting process and the values of *V*_*c*_ and *A*_*n*_. This is a consequence of the fact that (1 − Γ^*^/*C*) is negative for *C* < Γ^*^. For example, neglecting TPU, suppose that *W*_*c*_ < *W*_*j*_ for *C* < Γ^*^. In this case, min{*W*_*c*_, *W*_*j*_} = *W*_*c*_, so the min-*W* variant predicts Rubisco limitations with *V*_*c*_ = *W*_*c*_ and *A*_*n*_ = *A*_*c*_ (Figure **2a**). In contrast, the min-*A* variant uses the comparison *W*_*c*_ · (1 − Γ^*^/*C*) > *W*_*j*_ · (1 − Γ^*^/*C*), and thus predicts RuBP regeneration limitations with *V*_*c*_ = *W*_*j*_ and *A*_*n*_ = *A*_*j*_ (Figure **2b**), leading to a discrepancy where the min-*A* variant predicts a larger *V*_*c*_ and smaller *A*_*n*_ (Figure **2c**). From this analysis, we can see that the relationship between *A*_*c*_, *A*_*j*_, *A*_*p*_ and *A*_*n*_ in the min-*W* variant is

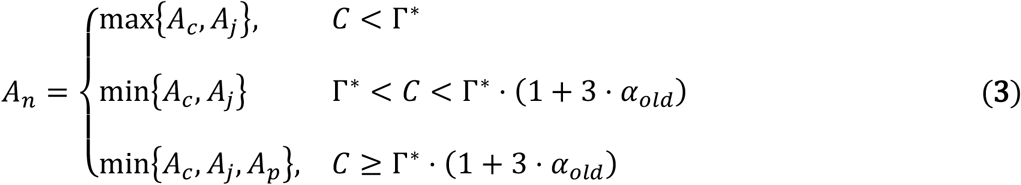

rather than Equation **2d**. Typical values of Γ^*^ are close to 40 μbar (approximately 40 μmol mol^-1^ or 40 ppm at typical atmospheric pressure), so this discrepancy between the variants only occurs at low values of *C*. Historically, such low values of *C* have been rarely accessed in gas exchange measurements because the available equipment could not easily reach them, but modern instruments can achieve lower values and researchers are now more likely to encounter *C* < Γ^*^. As will be shown below, this small difference prevents the min-*A* variant from matching measured CO_2_ response curves, and it can have a large influence on parameter estimation (Section **3.5**). Irrespective of any practical consequences, the min-*A* variant is not logically consistent because choosing the smallest net assimilation rate can lead to paradoxical results. For example, if *V*_*cmax*_ were to become zero when *C* < Γ^*^ due to substantial Rubisco deactivation (Section **3.4**), the min-*A* variant would nevertheless predict *V*_*c*_ = *W*_*j*_ and *A*_*n*_ = *A*_*j*_, as discussed above. Assuming a nonzero *J*, this is a contradictory prediction that *V*_*c*_ > 0 when *V*_*cmax*_ = 0. In contrast, choosing a minimal carboxylation rate always produces self-consistent results.
2. The two variants always disagree about the number of points where the rate-limiting process changes; these special values of *C* are called crossover points. In the min-*A* variant, *A*_*c*_ = *A*_*j*_ = −*R*_*L*_ when *C* = Γ^*^ (Equations **2a** and **2b**), so a crossover point always exists at *C* = Γ^*^ (Figure **2b**). However, in the min-*W* variant, *W*_*c*_ and *W*_*j*_ are not equal for *C* = Γ^*^ (Equations **1a** and **1b**), so a crossover point does not exist at *C* = Γ^*^ (Figure **2a**), except for the unlikely case that *J* happens to equal 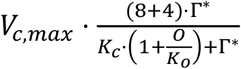 (Supplemental Section **S3.2**). In other words, there is a spurious crossover point at *C* = Γ^*^ in the min-*A* variant. This spurious crossover point has been noticed before (Dubois et al., 2007; Gu et al., 2010) and the FRL modification has been used to address this issue, although this modification does not fully reconcile the variants in all situations (Section **3.2**). Note that these discrepancies between the min-*W* and min-*A* approaches persist in the 3-*α* version of the FvCB model, where they appear for *C* < Γ^*^ ·(1 − *α*_*G*_ + 2 · *α*_*T*_) (Supplemental Section **S1.2**).

### 3.2. Rubisco Activity Does Not Always Limit Carboxylation at Low CO_2_ Concentrations

Some fitting tools that implement the min-*A* variant introduce a constraint in their code where *A*_*n*_ = *A*_*c*_ is enforced for *C* ≤ Γ^*^ (or possibly below another threshold value), even when Equation **2d** would predict otherwise (Figure **1** and Supplemental Section **S2.4**). Although rarely written explicitly as an equation, this is equivalent to replacing Equation **2d** with

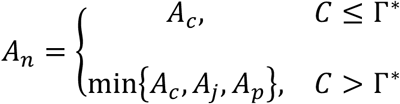

We refer to this as the “forced Rubisco limitation” (FRL) modification, and such variants as min-*A*+FRL variants. These variants agree with the min-*W* variant in situations where carboxylation is Rubisco limited for *C* ≤ Γ^*^.

However, the FRL modification will never be truly successful at fixing the min-*A* variant because there are no simple rules governing the progression of rate-limiting processes as *C* increases. Carboxylation can be limited by Rubisco activity or RuBP regeneration at low *C*, and a crossover between limitations is not guaranteed. Neglecting TPU, it is possible to analytically determine which process is limiting carboxylation at *C* = 0 and whether a crossover occurs (Supplemental Section **S3**). This process can be performed for many different values of *V*_*c,max*_ and *J*, producing a map that shows where each possible sequence of assimilation rates occurs in the min-*W* variant (Figure **3a**). When including TPU, it is more straightforward to simulate a CO_2_ response curve and extract the observed sequence of limitations; again, this can be performed for many values of *V*_*c,max*_ and *J*, producing a map (Figure **3b**).

**Figure 3:**
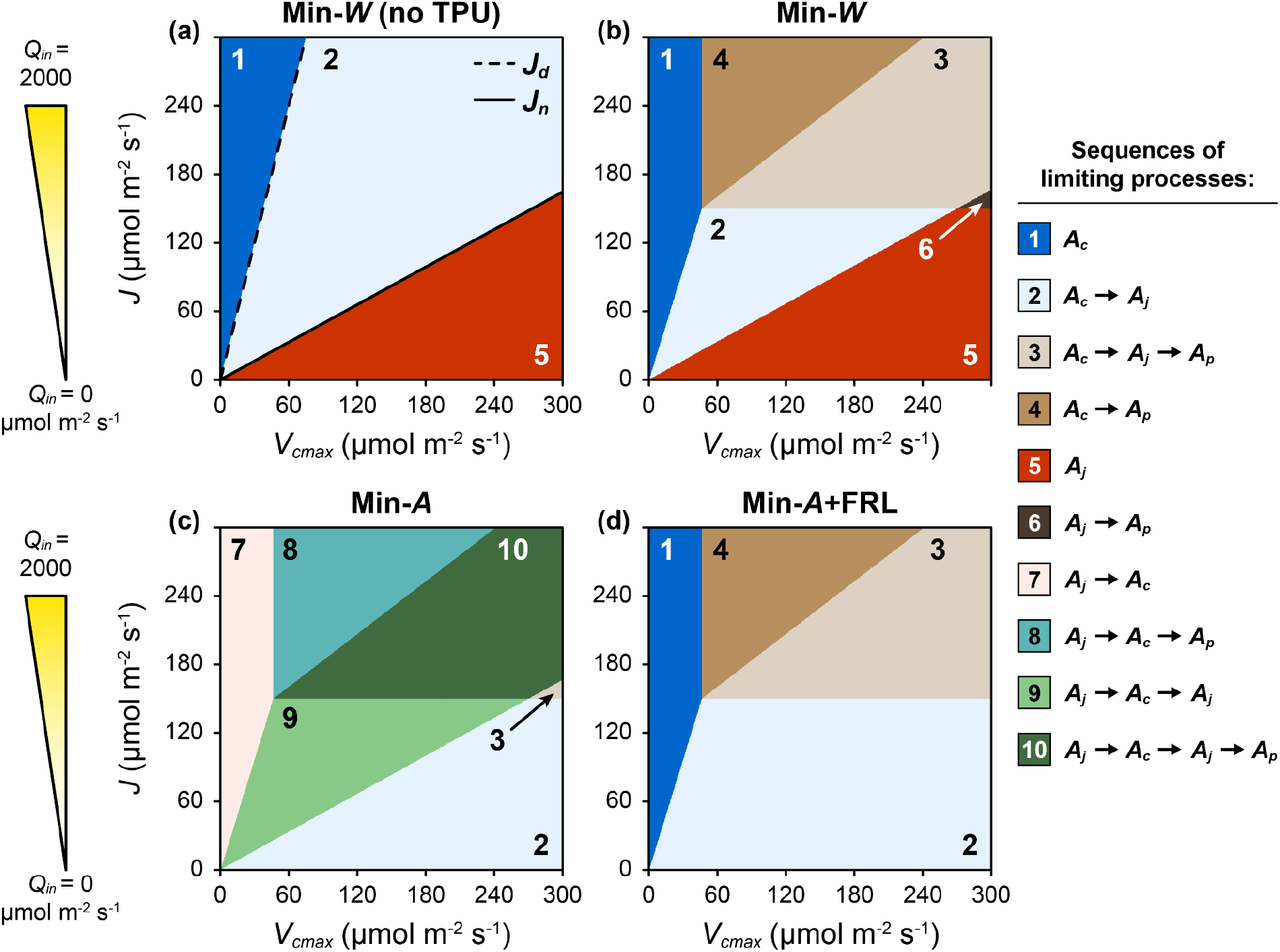
Predicted sequences of limiting processes as C increases from zero. (**a-d**) Maps of (V_c,max_, J)-space, where labels within each region indicate the sequences as calculated by the min-W variant (neglecting or including TPU) (**a-b**), the min-A variant (**c**), or the min-A+FRL variant (**d**). The triangles to the left of (**a**) and (**c**) indicate an approximate relation between J and Q_in_. Dashed and solid black lines in (**a**) show J_d_ and J_n_ (Equations **A6** and **A7** in Supplemental Section **S3**). All labels in (**b-d**) refer to observed sequences below a threshold of 2000 μbar. All parameters other than V_c,max_ and J were set to the values specified in Table **1**.

This analysis shows that when *V*_*c,max*_ is relatively small compared to *J*, the min-*W* variant predicts Rubisco limitations for all values of *C* (dark blue regions labeled “1” in Figure **3**). This has been observed in modified tobacco with reduced Rubisco content (von Caemmerer et al., 1994), and this situation could potentially occur in plants under nitrogen stress (leading to low *V*_*c,max*_) or in leaves exposed to high incident light levels (leading to high *J*). Likewise, when *J* is relatively small compared to *V*_*c,max*_, the min-*W* variant predicts RuBP regeneration limitations for all values of *C* (red regions labeled “5” in Figure **3**). This situation could potentially occur in leaves exposed to low incident light levels (leading to low *J*), although this may be difficult to observe in practice due to a concurrent reduction in *V*_*c,max*_ that tends to occur at low *Q*_*in*_ (Lochocki et al., 2025; Sage et al., 2002; Taylor et al., 2022). Actual values of *Q*_*in*_ corresponding to high or low *J* depend strongly on details of leaf biochemistry and growth environment, but for many plants, *Q*_*in*_ ≤ 100 μmol m^-2^ s^-1^ would correspond to low *J* while *Q*_*in*_≥ 1800 μmol m^-2^ s^-1^ would correspond to high *J*. The maps also show that *A*_*c*_ → *A*_*j*_ → *A*_*p*_ is not the only predicted sequence involving TPU limitations. In fact, a transition directly from Rubisco limited to TPU limited carboxylation (*A*_*c*_ → *A*_*p*_) can be observed in plants grown in low-light conditions (Sharkey, 2019). Note that there is no region corresponding to *A*_*j*_ → *A*_*c*_, indicating that this transition is not predicted to occur for typical parameter values (Supplemental Sections **S3** and **S4**).

Similar maps generated using the min-*A* and min-*A*+FRL variants show large differences relative to the min-*W* map (Figures **3c** and **3d**). Because of the spurious crossover point at Γ^*^ in the min-*A* variant (Section **3.1**), each region of the min-*A* map has one extra step compared to its min-*W* analog, several of which do not occur in the min-*W* map. For example, the *A*_*c*_ → *A*_*p*_ region of the min-*W* map (dark tan region labeled “4” in Figure **3b**) becomes an *A*_*j*_ → *A*_*c*_ → *A*_*p*_ region in the min-*A* map (teal region labeled “8” in Figure **3c**). Because the min-*A*+FRL variant enforces Rubisco-limited assimilation at low *C*, its map lacks several sequences that are present in the min-*W* map. For example, the *A*_*j*_ region of the min-*W* map becomes an *A*_*c*_ → *A*_*j*_ region in the min-*A*+FRL map. These results demonstrate that the min-*W* and min-*A* variants do not predict the same transitions between limiting processes, and that the FRL modification does not fully bring the min-*A* variant into agreement with the min-*W* variant.

### 3.3. TPU Can Only Limit Carboxylation When Photosynthesis and Photorespiration Are Net Consumers of Free Inorganic Phosphate

Additional differences between the min-*W* and min-*A* variants can appear when *α*_*old*_ > 0. This parameter is related to TPU and appears in Equations **1c** and **2c**. In the min-*W* variant, nonzero *α*_*old*_ introduces a downward slope to *W*_*p*_ (Figure **4a**). The resulting decrease in *A*_*p*_ with increasing *C*, termed “reverse sensitivity,” is an identifying signature of TPU limitations, and the inclusion of *α*_*old*_ is essential to fit CO_2_ response curves exhibiting reverse sensitivity (Busch et al., 2018; Harley and Sharkey, 1991). In the min-*A* variant, nonzero *α*_*old*_ similarly introduces a downward slope to *A*_*p*_ at high values of *C*, but also produces divergent behavior at lower values of *C* (Figure **4b**). Mathematically, this behavior occurs because the denominator of Equation **2c** becomes zero when *C* = Γ^*^ · (1 + 3 · *α*_*old*_), and *A*_*p*_ approaches −∞ as *C* approaches Γ^*^ · (1 + 3 · *α*_*old*_) from below. This issue does not occur for *W*_*p*_ because Equation **1c** sets *W*_*p*_ to ∞ for *C* ≤ Γ^*^ · (1 + 3 · *α*_*old*_); however, it will occur if this condition is omitted from Equation **1c**.

**Figure 4:**
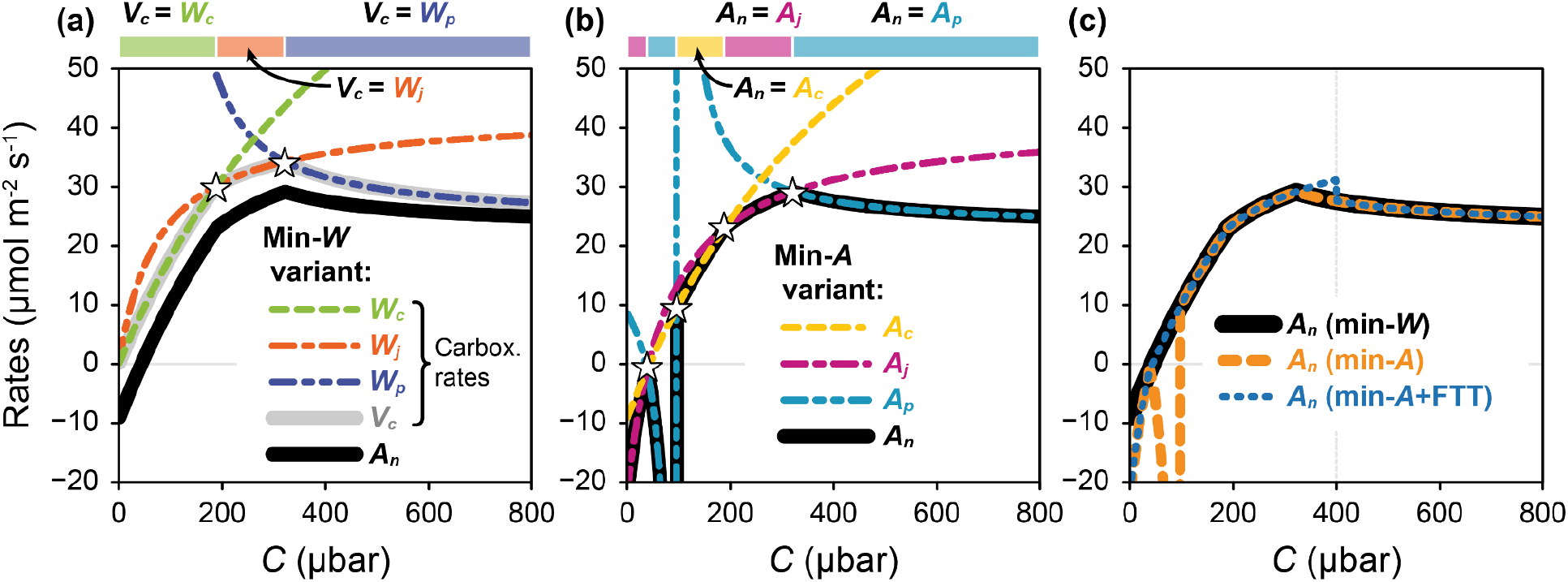
Comparison of min-W, min-A, and min-A+FTT variant outputs for identical input parameter values with α_old_ > 0. (**a**,**b**)A_n_ (thick solid lines) as determined from potential carboxylation or assimilation rates (dashed lines) using Equations **1** and **2**, respectively. Crossover points where the rate-limiting process changes are marked with white stars. Solid rectangles indicate ranges of C where carboxylation or assimilation is limited by Rubisco, RuBP regeneration, or TPU. (**c**) Direct comparison of the variants across a range of C values. The following parameter values were used for these calculations: V_c,max_ = 120 μmol m^-2^ s^-1^,T_p_ = 8 μmol m^-2^ s^-1^, and α_old_ = 0.5. All other parameters were set to the values specified in Table **1**.

Some fitting tools introduce a constraint in their code where TPU cannot set the net CO_2_ assimilation rate below a fixed threshold value of *C*, which is often chosen to be 400 µbar (Supplemental Section **S2.4**). Although rarely written explicitly as an equation, when using the min-*A* variant this is equivalent to replacing Equation **2c** with

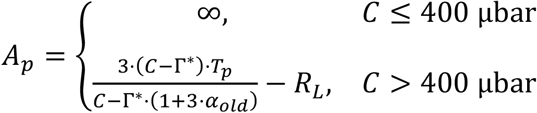

or when using the min-*W* variant, replacing Equation **1c** with

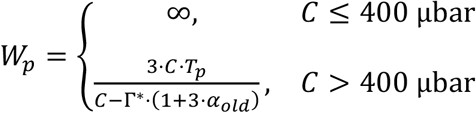

We refer to this as the “fixed TPU threshold” (FTT) modification, and such variants as the min-*A*+FTT or min-*W*+FTT variants. (Note that the min-*A*+FTT and min-*A*+FRL variants can be combined to form another variant with both types of modifications: the min-*A*+FRL+FTT variant.) These FTT variants agree with the min-*W* variant in situations where *W*_*p*_ is not the smallest carboxylation rate below the threshold value of *C*. However, if the min-*W* variant predicts *A*_*n*_ = *A*_*p*_ for Γ^*^ · (1 + 3 · *α*_*old*_) < *C* < 400 μbar, then the min-*W*+FTT and min-*A*+FTT variants will predict *A*_*n*_ ≠ *A*_*p*_ in that range, and then suddenly drop down to *A*_*n*_ = *A*_*p*_ above the threshold (Figure **4c**). TPU limitations below 400 μbar are rare, but have been observed (Harley and Sharkey, 1991). Another potential issue with the FTT modification is that the divergent behavior of *A*_*p*_ will persist if the fixed threshold value is smaller than Γ^*^ ·(1 + 3 · *α*_*old*_).

The FTT modification will never be truly successful at fixing the issues with Equation **2c** because it does not account for an important biochemical aspect of TPU limitations: TPU can only limit carboxylation when the net effect of photosynthesis and photorespiration is to shrink the pool of free inorganic phosphate (P_i_) in the chloroplast by incorporating it into organic compounds, a process referred to as P_i_ consumption. P_i_ is required to synthesize adenosine triphosphate (ATP) via photophosphorylation, which in turn is used to regenerate RuBP. To achieve sustained ATP synthesis and RuBP regeneration, the net rate of P_i_ consumption due to photosynthesis and photorespiration (*R*_*pc*_) must be balanced by the rate at which P_i_ is returned to the chloroplast through the utilization of triose phosphate for sugar production (*T*_*p*_); otherwise, the pool of P_i_ would become depleted (Harley and Sharkey, 1991; Sharkey, 1985; von Caemmerer, 2000). In other words,

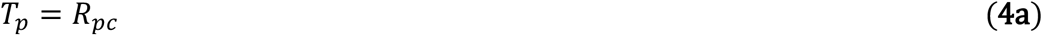

when TPU limits carboxylation. This requirement can be used to derive Equation **1c**, where the key step in the derivation is to relate *R*_*pc*_ to the RuBP carboxylation rate. Here we show that the restriction *C* > Γ^*^ · (1 + 3 · *α*_*old*_) is a natural consequence of this derivation.

By considering the stoichiometry of the carboxylation and oxygenation cycles and allowing for an additional release of P_i_ during photorespiration due to glycolate carbon that remains in the cytosol, it can be shown that

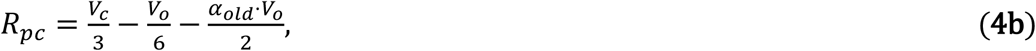

where *V*_*o*_ is the rate of RuBP oxygenation (Harley and Sharkey, 1991; von Caemmerer, 2000). Next, it can also be shown that *V*_*o*_ is related to *V*_*c*_ via *V*_*o*_ = 2 · Γ^*^ · *V*_*c*_/*C* (von Caemmerer, 2000), allowing us to eliminate *V*_*o*_ from Equation **4b**:

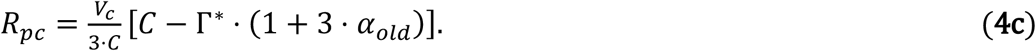

Here it is essential to note that, first, when *C* = Γ^*^ · (1 + 3 · *α*_*old*_), the net rate of P_i_ consumption is zero. Second, for smaller values of *C, R*_*pc*_ is negative, indicating that photosynthesis and photorespiration are actually releasing P_i_. In both of these situations, it is not possible for TPU to become a limiting factor because photosynthesis and photorespiration are not shrinking the pool of P_i_ in the chloroplast, and hence the return of P_i_ via TPU is not necessary to prevent P_i_ depletion.

Finally, when TPU limits the carboxylation rate, *V*_*c*_ is denoted by *W*_*p*_; in this case, we can now use Equations **4a** and **4c** to express the requirement for TPU limitation as

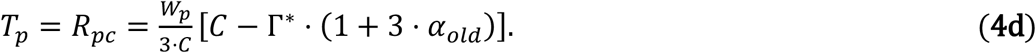

Solving Equation **4d** for *W*_*p*_ and remembering that photosynthesis and photorespiration only consume P_i_ when *C* > Γ^*^ · (1 + 3 · *α*_*old*_) yields Equation **1c**. This lower *C* threshold for TPU limitations is a consequence of leaf biochemistry and its inclusion is guaranteed to prevent numerical errors such as negative or undefined *W*_*p*_, unlike the fixed threshold used in the FTT modification. Although this threshold could be included in Equation **2c** from the min-*A* variant as a conditional statement like the one in Equation **1c**, none of the studies in the literature survey (Supplemental Section **2**) take this approach. Furthermore, doing so would not address the other issues with the min-*A* variant (Sections **3.2, 3.3**, and **3.5**). Note that in the 3-*α* version of the model, the biochemically derived *C* threshold for TPU limitations becomes *Γ*^*^ ·(1 − *α*_*G*_ + 2 · *α*_*T*_) · (1 + 3 · *α*_*G*_ + 4 · *α*_*S*_ + 6 · *α*_*T*_) (Supplemental Section **S1.2**).

### 3.4. The Min-W Variant Can Be Extended To Include Substantial Rubisco Deactivation and RuBP Depletion at Low CO_2_ Concentrations but the Min-A Variant Cannot

Besides the processes explicitly included in Equations **1** and **2**, Rubisco deactivation and RuBP depletion may also influence net CO_2_ assimilation rates. Rubisco deactivation can occur when a leaf is held under low CO_2_ conditions or under low light, likely by increasing the fraction of decarbamylated Rubisco, and the result of this process can be modeled as a reduction in *V*_*cmax*_ (Lochocki et al., 2025; Taylor et al., 2022; Sage et al., 2002). The min-*W* and min-*A* variants make different predictions in the hypothetical extreme case of full Rubisco deactivation for *C* < Γ^*^, where *V*_*cmax*_ would become zero. Assuming no other parameter values change, *W*_*c*_ would also become zero but *W*_*j*_ and *W*_*p*_ would remain positive, so the min-*W* variant would predict Rubisco limitations with *A*_*n*_ = *A*_*c*_ = −*R*_*L*_ (Equation **1**). Yet, *A*_*j*_ would be less than *A*_*c*_ = −*R*_*L*_, so the min-*A* variant would predict a different limiting process and a smaller value of *A*_*n*_ (Equation **2**). In other words, the min-*A* variant does not allow substantial Rubisco deactivation to limit net CO_2_ assimilation at low *C*, even when no Rubisco sites are active.

Similarly, the min-*A* variant does not predict a change in *A*_*n*_ due to substantial RuBP depletion, which can occur when *C* < Γ^*^ because RuBP cannot be regenerated in this range. RuBP regeneration depends not only on electron transport and the availability of P_i_, but also on the supply of 3-phosphoglyceric acid (PGA) from photosynthesis and photorespiration, which is reduced to unbound RuBP in a multi-step enzymatic pathway (von Caemmerer, 2000). When *C* < Γ^*^, the supply of carbon as PGA is insufficient to regenerate enough RuBP, eventually depleting the pool of unbound RuBP. This can be understood by considering the carbon stoichiometry of the reduction pathway, where Rubisco uses RuBP at a rate of *V*_*c*_ + *V*_*o*_ and PGA is produced at a rate of 2*V*_*c*_ + 1.5*V*_*o*_ (von Caemmerer, 2000). RuBP and PGA are five- and three-carbon molecules, respectively, so the difference between carbon supply and demand (Δ_*c*_) is

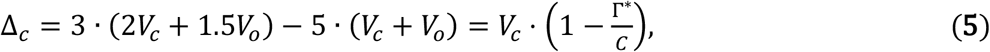

where *V*_*o*_ = 2 · Γ^*^ · *V*_*c*_/*C* has been used to simplify the final expression. There is a carbon surplus (Δ_*c*_ > 0) for *C* > Γ^*^, which is exported from the chloroplast as triose phosphates. Carbon supply and demand are exactly equal (Δ_*c*_ = 0) when *C* = Γ^*^, and there is a deficit (Δ_*c*_ < 0) for *C* < Γ^*^. In the latter case, continued Rubisco activity draws from the chloroplastic pool of unbound RuBP without replenishing it.

Modeling RuBP depletion caused by insufficient carbon supply is more complex than Rubisco deactivation because it does not simply alter the value of a parameter such as *V*_*cmax*_ or *J*.

Although not traditionally considered as part of the FvCB model, here we include it in the min-*W* variant by adding a fourth potential limitation to Equation **1d**:

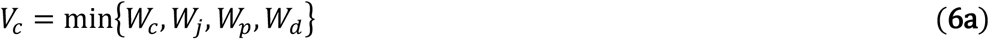

where *W*_*d*_ is the carboxylation rate limited by substantial RuBP depletion. *W*_*d*_ is taken to be zero when *C* has been held below Γ^*^ long enough for Rubisco activity to completely deplete the pool of unbound RuBP, and is otherwise assumed to not limit carboxylation. This can be described mathematically as

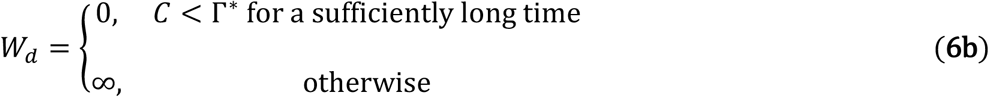

Equation **6b** is not precise and omits two key details – incomplete RuBP depletion can lead to a small but nonzero *W*_*d*_ at low *C*, and *W*_*d*_ should depend continuously on *C*, RuBP pool size, and time. Nevertheless, this simplistic equation is instructive for understanding long-term steady-state net CO_2_ assimilation at low *C*. When substantial RuBP depletion limits carboxylation, *V*_*c*_ = *W*_*d*_ and the corresponding net CO_2_ assimilation rate (*A*_*d*_) can be found with Equations **1a** and **6b**:

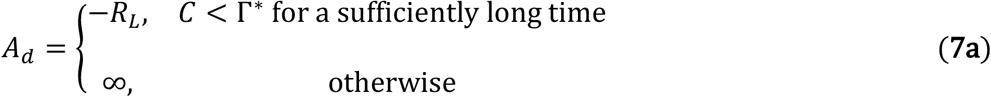

With this, RuBP depletion can also be incorporated into the min-*A* variant by adding a fourth potential assimilation rate to Equation **2d**:

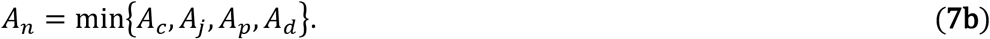

Equations **6** and **7** make contradictory predictions for *A*_*n*_ when *C* is below Γ^*^ for enough time to ensure substantial RuBP depletion. Assuming otherwise typical model parameter values in this scenario, *W*_*c*_, *W*_*j*_, and *W*_*p*_ would each be positive, so *W*_*d*_ = 0 would always be the smallest carboxylation rate, and the min-*W* variant would predict RuBP depletion limitations with *A*_*n*_ =*A*_*d*_ = −*R*_*L*_ (Equation **6**). Conversely, *A*_*c*_ and *A*_*j*_ would each be smaller than −*R*_*L*_, so the min-*A* variant would never predict *A*_*n*_ = *A*_*d*_. In fact, Equation **7** would always predict the same rate and limiting process as Equation **2**. In other words, the min-*W* variant predicts a shift of *A*_*n*_ to *A*_*d*_ = −*R*_*L*_ when substantial RuBP depletion occurs at low CO_2_ concentrations, while the min-*A* variant does not allow substantial RuBP depletion to limit net CO_2_ assimilation.

### 3.5. Comparing Both Variants to Measured CO_2_ Response Curves

Since the min-*W* and min-*A* variants make different predictions for *C* < Γ^*^, even when considering substantial Rubisco deactivation and RuBP depletion, it is possible to test them against measurements to determine which variant better represents reality. To do this, thirty-six CO_2_ response curves were measured from *Nicotiana tabacum* (tobacco) cv. Samsun leaves at multiple incident light intensities using Licor LI-6800 portable gas exchange systems (Supplemental Section **S5.1**). Reference CO_2_ concentration set-points ranging from 10 − 1800 μmol mol^-1^ were used, where the lowest values ensured several points in each curve with *C* < Γ^*^. Each curve was fit with the *PhotoGEA* R package (Lochocki, 2025; Lochocki et al., 2025) on a *C*_*i*_ basis (i.e., by setting *C* = *C*_*i*_) using tobacco temperature response parameters (Sharkey et al., 2007) and allowing *V*_*c,max*_, *J, R*_*L*_, *T*_*p*_, and *α*_*old*_ to vary. *PhotoGEA* uses Equation **1** by default, but can optionally use Equations **2** or **6** instead, and can also optionally apply the FRL or FTT modifications. The fits were performed in four ways:

- min-*W variant (all C*_*i*_): The entire curve was fit twice using Equations **1** and **6**, and the result with smaller root mean square error (RMSE) was chosen as the best fit. If Equation **6** produced the best fit, the curve was considered to exhibit *A*_*d*_ limitations; otherwise, the curve was not considered to exhibit *A*_*d*_ limitations.
- min-*A+FTT variant (all C*_*i*_): The entire curve was fit using Equation **2** with a fixed TPU threshold of 400 μbar. There is no need for an additional fit using Equation **7** because Equations **2** and **7** make identical predictions.
- min-*W variant (C*_*i*_ *> 45 μbar)*: Points from the curve where *C*_*i*_ is above 45 µbar were fit using Equation **1**. Requiring *C*_*i*_ > 45 μbar ensured that no points with *C* < Γ^*^ were used for the fit, since tobacco Γ^*^ is approximately 39 μbar at the measurement temperature of 27 °C.
- min-*A+FTT variant (C*_*i*_ *> 45 μbar)*: Points from the curve where *C*_*i*_ is above 45 μbar were fit using Equation **2** with a fixed TPU threshold of 400 μbar.

Note that the min-*W* variant predicts *A*_*n*_ = −*R*_*L*_ when either substantial Rubisco deactivation or RuBP depletion limits net CO_2_ assimilation for *C* < Γ^*^ (Section **3.4**), so it is not possible to unambiguously attribute an observed *A*_*d*_ limitation to either process. Because of this, we consider *W*_*d*_ and *A*_*d*_ to represent both processes in the context of curve fitting. Also note that the min-*A*+FTT variant is used for these comparisons because the divergent behavior of the min-*A* variant with nonzero *α*_*old*_ severely interferes with the fitting process (Section **3.3**).

For a curve without such *A*_*d*_ limitations, an extrapolation of the fit made using the min-*W* variant for points where *C*_*i*_ is above 45 µbar agrees well with the measured points at lower *C*_*i*_ (Figure **5a**), suggesting that a good fit across all measured points could be achieved using Equation **1**. However, an extrapolation of the fit made using the min-*A*+FTT variant greatly diverges from the measured assimilation rates for *C*_*i*_ below 45 µbar, even though the two fits are identical for higher *C*_*i*_ (Figure **5b**). For a curve with *A*_*d*_ limitations, a similar extrapolation of the min-*W* fit made using points where *C*_*i*_ is above 45 µbar does not match the measured points at lower *C*_*i*_, but *A*_*d*_ = −*R*_*L*_ does lie close to those points (Figure **5e**), suggesting that a good fit across all measured points could be achieved using Equation **6**. An extrapolation of the min-*A*+FTT fit is even further from the measured points where *C*_*i*_ is below 45 µbar (Figure **5f**), but the min-*A* variant never predicts *A*_*n*_ = *A*_*d*_, so a good fit across all *C*_*i*_ is unlikely.

**Figure 5:**
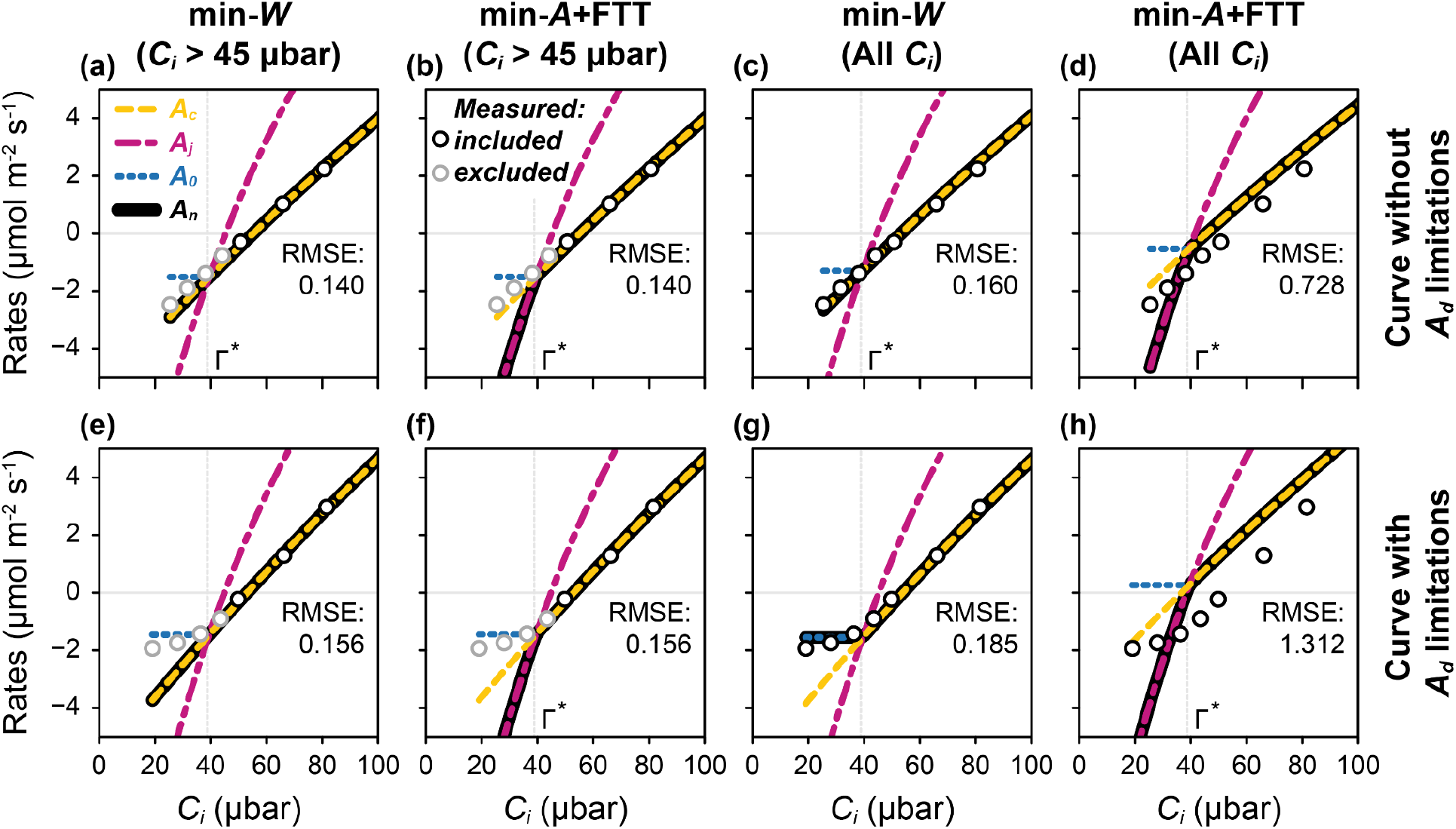
Testing FvCB model variants against measured CO_2_ response curves. Fits are shown for one curve without A_d_ limitations (**a-d**) and one with A_d_ limitations (**e-h**), both of which were measured with Q_in_ = 600 µmol m^-2^ s^-1^. Fits were made using the min-W (**a**,**c**,**e**,**g**) or min-A+FTT variant (**b**,**d**,**f**,**h**), either limited to C_i_ above 45 μbar (**a**,**b**,**e**,**f**) or using all measured points (**c**,**d**,**g**,**h**). In each panel, measured points that were included in or excluded from the fit are shown as open black and gray circles, respectively. RMSE values are calculated from only the points included in each fit. A zoomed-in portion of each curve is shown to highlight differences at low C_i_; see Supplemental Section **S5** for expanded views showing the fits across the entire range of measured C_i_.

As expected from these extrapolations, the min-*W* variant is able to closely match the measured points in each curve when fitting the entire range of *C*_*i*_ (Figures **5c** and **5g**), while the min-*A*+FTT variant is not (Figures **5d** and **5h**). This is reflected in the RMSE values, where a lower RMSE indicates a better fit. The RMSE values of the min-*W* fits barely increase when adding the points with *C*_*i*_ < 45 μbar (0.140 vs. 0.160 for the curve without *A*_*d*_ limitations and 0.156 vs. 0.185 for the curve with *A*_*d*_ limitations), while there is a large increase in the min-*A*+FTT RMSE values (0.140 vs. 0.728 for the curve without *A*_*d*_ limitations and 0.156 vs. 1.312 for the curve with *A*_*d*_ limitations) (Figure **5**).

These results hold across the entire set of thirty-six curves, where eighteen were found to exhibit *A*_*d*_ limitations at low *C*_*i*_ (Figure **6**). When fitting points at all *C*_*i*_, the min-*A*+FTT variant generally produces larger RMSE values than the min-*W* variant, especially for curves that exhibit *A*_*d*_ limitations (Figure **6a**). Across all curves, the mean RMSE for the min-*W* fits is 0.32 μmol m^-2^ s^-1^, while the mean RMSE for the min-*A*+FTT fits is more than twice as large at 0.79 μmol m^-2^ s^-1^ (see also Supplemental Figure **S10**). Fits made with the min-*W* variant have similar RMSE values when including or excluding the points where *C*_*i*_ is below 45 µbar, but fits made with the min-*A*+FTT variant are appreciably worse when including points where *C*_*i*_ is below 45 μbar, even for curves that do not exhibit *A*_*d*_ limitations. Overall, this indicates that the min-*W* variant is able to represent leaf net CO_2_ assimilation across all *C*_*i*_ values when potential limitations due to Rubisco deactivation and RuBP depletion are considered for *C* < Γ^*^ (Equation **6**), while the min-*A*+FTT variant cannot.

**Figure 6:**
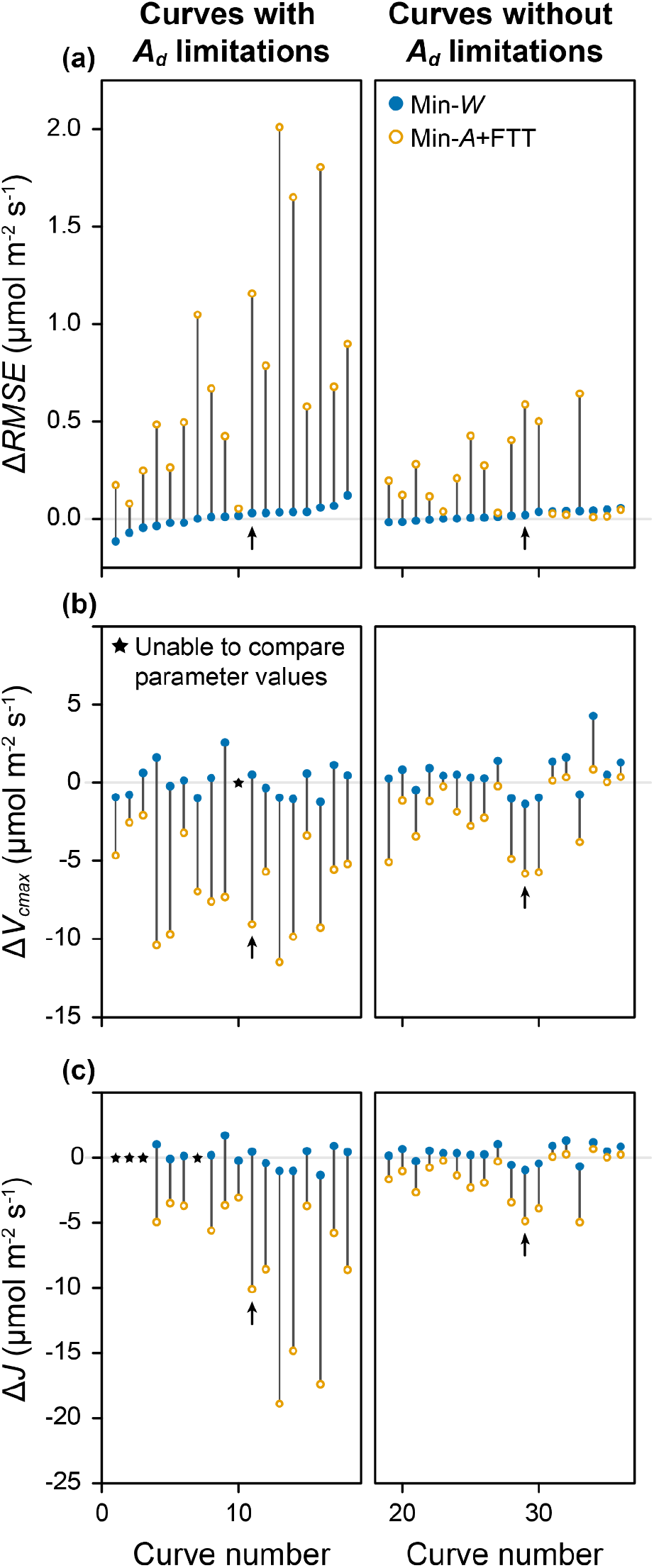
Comparing fit results from thirty-six CO_2_ response curves. Differences in (**a**) RMSE, (**b**) V_cmax_, and (**c**) J that occur when including all points in the fit as compared to only using points where C_i_ is above 45 µbar, where 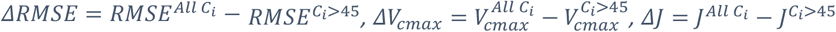, and superscripts indicate which points were fit. Values from min-W and min-A+FTT variant fits are shown as filled blue circles and open yellow circles, respectively. Vertical gray lines connect the values from each variant for each curve. Results from curves that exhibit or do not exhibit A_d_ limitations are shown in the left and right columns, respectively. Some parameter values could not be reliably estimated from some curves; such parameters are marked with black stars. The two curves shown in Figure **5** are marked with black arrows.

Because the min-*A*+FTT variant does not closely match the observed values of *A*_*n*_ for *C* < Γ^*^, parameter values estimated using this variant are greatly influenced by including points where*C*_*i*_ is below 45 µbar (Figures **6b-c**). As compared to fits made with the min-*W* variant, the min-*A*+FTT variant tends to underestimate both *V*_*cmax*_ and *J*. Values of *V*_*cmax*_ from min-*A*+FTT fits can be as much as 12 μmol m^-2^ s^-1^ lower (as much as 23% lower on a relative basis), while values of *J* can be as much as 17 µmol m^-2^ s^-1^ lower (as much as 12% lower on a relative basis).

## 4. Discussion

The broad goal of this study was to summarize the most commonly used variants of the FvCB model and to compare them using mathematical and experimental approaches, providing what we hope is a useful resource for those learning the history of the development of this model.

We also uncovered some issues with several of these variants and indicated the problems they can pose for FvCB model results and their interpretation. First, we showed that the min-*W* and the min-*A* variants disagree about the value of *A*_*n*_ and the rate-limiting process whenever *C* < Γ^*^ (Section **3.1**). The FRL modification can address this discrepancy in some situations but does not fully reconcile the two variants (Section **3.2**). Second, we demonstrated that unrealistic predictions can occur when the biochemically-derived lower *C* threshold for TPU limitations, Γ^*^ · (1 + 3 · *α*_*old*_), is not included in the equations, or when a fixed threshold is used instead (Section **3.3**). Third, we showed that substantial Rubisco deactivation and RuBP depletion can be included in the min-*W* variant, but that the min-*A* variant cannot easily accommodate these processes for *C* < Γ^*^ (Section **3.4**). Finally, we demonstrated that these issues prevent the min-*A*+FTT variant from closely fitting measured CO_2_ response curves that include points where *C* < Γ^*^, a potential source of error when estimating FvCB parameter values (Section **3.5**).

Half of the curves fit in this study exhibited signs of substantial Rubisco deactivation or RuBP depletion for *C* < Γ^*^, requiring Equation **6** rather than Equation **1** to achieve a good fit. Because the min-*A* variant never allows these processes to limit assimilation at low *C*, it was unable to produce good fits to all of the measured curves. At present, it is not possible to predict whether a particular leaf will exhibit *A*_*d*_ limitations, making Equations **1** and **6** useful for describing curves but not necessarily for simulating assimilation rates when *C* < Γ^*^. For typical *A*-*C*_*i*_ curves, where fewer points are recorded at very low CO_2_ concentrations, Equation **6** may not be needed at all. The treatment of Rubisco deactivation and RuBP depletion in Equation **6** is simplistic and could be improved, but the main intent here is to demonstrate that it is straightforward to include these processes in the min-*W* variant but not in the min-*A* variant. A more realistic approach would be to include steady state or time dependent RuBP pool sizes,where the latter is included in *e*-photosynthesis (Zhu et al., 2007, 2013), a dynamic photosynthesis model, but would not be compatible with the steady-state FvCB model.

Parameter estimation from experimentally measured CO_2_ response curves is a major application of the FvCB model. Whenever a curve fitting tool uses the min-*A*, min-*A*+FRL, or min-*A*+FTT variants rather than the min-*W* variant, there is a potential for errors in the estimated parameter values. These errors can occur whenever any of the points along the curve are measured in conditions where the min-*W* variant disagrees with the other variants. This is likely to happen in the following situations:

- If the curve extends to low CO_2_ concentrations (*C* roughly below 40 μbar), some of the measured points may satisfy *C* < Γ^*^, and the min-*A* variant will assign the wrong limiting process in this range (Section **3.1**). This issue persists even when considering substantial Rubisco deactivation and RuBP depletion (Section **3.4**).
- If the curve extends to low CO_2_ concentrations and was measured with low light levels around *Q*_*in*_ = 100 μmol m^-2^ s^-1^ (such that assimilation is limited by RuBP regeneration across the entire measured range), then the min-*A* variant will predict a sensitivity of the fit to *V*_*c,max*_, even though *V*_*c,max*_ would have no influence on the fit using the min-*W* variant. This error will persist even when adding the FRL modification (Section **3.2**).
- If the curve exhibits strong TPU limitations with reverse sensitivity, a nonzero value of *α*_*old*_ will be necessary. This will cause divergent behavior in the min-*A* variant at lower CO_2_ concentrations, preventing a good fit (Section **3.3**).
- If the curve exhibits strong TPU limitations where *A*_*n*_ = *A*_*p*_ for *C* below 400 μbar, the min-*A*+FTT variant will not be able to represent this behavior (Section **3.3**).

Here we show that estimates of *J* and *V*_*c,max*_ made using the min-*A*+FTT variant are typically lower than those from the min-*W* variant, with underestimates exceeding 20%. Because the differences between the variants only appear for certain environmental conditions and plant characteristics, the only way to assess errors introduced by the min-*A* variant for a particular CO_2_ response curve is to compare parameter estimates against those made using the min-*W* variant. Rather than attempting to quantify errors in a range of situations, it is simpler to always use the min-*W* variant.

Across the scientific literature, most CO_2_ response curves are measured under high light and may contain only one or two points at very low CO_2_ concentrations. It is also rare for TPU to be present for *C* below 400 µbar. Thus, the min-*A* variant is not likely to have caused significant errors in published work. It is also possible that some researchers intentionally avoid the complexities of fitting curves with points where *C* < Γ^*^, either by only measuring points at higher CO_2_ set-points, or by excluding potentially problematic points (e.g., where *C*_*i*_ < Γ^*^ or *A*_*n*_ < 0) when fitting. Often this practice is not documented, even in recent comprehensive guides to photosynthetic gas exchange measurements such as Busch et al. (2024), but it is likely common enough to mitigate problems caused by the min-*A* approach. Nonetheless, the min-*A* variant has no benefits over the min-*W* variant and is equally complex, leaving little reason for its use. When it is desired to express the model equations using only assimilation rates, the form presented in Equation **3** could be used instead.

As an argument against using the min-*A* variant despite the lack of current impact, the likelihood of encountering some of these problematic situations may become larger as research trends change and new research areas are formed. For example, a recently developed method for estimating leaf cuticle conductance using gas exchange measurements actually necessitates the measurement of CO_2_ response curves under low irradiance (Márquez et al., 2022), a situation where the min-*A* and min-*A*+FRL variants can produce incorrect parameter estimates. The Laisk method for estimating 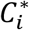 and *R*_*L*_ also requires low-light CO_2_ response curves (Laisk, 1977; Walker and Ort, 2015). Efforts to improve crop yield and food security by engineering plants with new Rubisco homologs may create situations where Rubisco-related parameters like Γ^*^ and *K*_*c*_ take values significantly different from the ones in Table **1** (Amaral et al., 2024; Carmo-Silva et al., 2015; Long et al., 2015; Parry et al., 2013). The most extreme situations may occur in the field of astrobiology, where researchers study photosynthesis in hypothetical extraterrestrial conditions with ambient gas environments and biochemical parameters that may be very different from those found on Earth (Covone et al., 2021; Lehmer et al., 2021; Lingam and Loeb, 2019; Mullan and Bais, 2018). It is therefore more important than ever for plant scientists to clearly describe their curve fitting methods, including the equations used (either min-*W* or min-*A*), whether any measured points satisfied *C* < Γ^*^, and whether any points at low *C*_*i*_ or *A*_*n*_ were excluded from fits.

A literature survey indicates that the min-*W* and min-*A* approaches have been used with roughly equal frequency. Yet, the summarization and comparison of FvCB model variants presented here indicates that the min-*W* approach, along with the biochemically-derived lower threshold for TPU limitations, is the current state of the art, and likely the most appropriate choice in many cases. Among min-*W* variants, the primary decision for researchers is to determine whether a situation calls for a version of the FvCB model that uses *α*_*old*_ (Equation **1**, the 1-*α*+BTT+min-*W*+NFL variant) or one that accounts for separate glycolate pathways via *α*_*G*_, *α*_*S*_, and *α*_*T*_ (Supplementary Equation **S1**, the 3-*α*+BTT+min-*W*+NFL variant).

## Supporting information

Supplemental Sections S1-S5, including Supplemental Figures S1-S10

## Acknowledgements

The authors thank Stephen P. Long, Carl J. Bernacchi, Coralie E. Salesse-Smith, Yi Xiao, and Scott Rohde for helpful discussions.

This work was supported by Bill & Melinda Gates Agricultural Innovations grant investment ID 57248.

Any opinions, findings, and conclusions or recommendations expressed in this publication are those of the authors and do not necessarily reflect the views of the US Department of Agriculture. Mention of trade names or commercial products in this publication is solely for the purpose of providing specific information and does not imply recommendation or endorsement by the US Department of Agriculture. USDA is an equal opportunity provider and employer.

## Author Contributions

E.B.L. contributed to the conceptualization, modeling, data analysis and writing (original draft).

J.M.M. contributed to conceptualization and revising the manuscript.

## Data Availability

All data and R scripts for reproducing all analysis contained in this work are available online at https://github.com/ripeproject/FvCB-min-A.

## Supporting Information

The supporting information includes Section **S1** (*Variations in FvCB Model Equations and Nomenclature*), Section **S2** (*Literature Survey*), Section **S3** (*CO*_*2*_ *Response in the Min-W Variant*) Section **S4** (*Sequences of Limiting Processes for High Γ*^*^ *and/or Low K*_*c*_), and Section **S5** (*Fitting Experimentally Measured CO*_*2*_ *Response Curves*).

