## Supplemental Sections S1-S5, including Supplemental Figures S1-S10 for "Widely Used Variants of the Farquhar-von-Caemmerer-Berry Model Can Cause Errors in Parameter Estimation"

### Supplementary Information

### S1. Variations in FvCB Model Equations and Nomenclature

#### S1.1 Non-Essential Variations

A large number of variations in FvCB model equations and nomenclature can be found in the literature. Most of them do not affect the general conclusions about the variants presented here; the essential differences are described in Table **2** of the main text. The variations described in the following (non-exhaustive) list are considered non-essential for our purposes:

- Strictly speaking, $C$ should be the partial pressure of $CO_2$ in the chloroplast ($C_c$), where Rubisco is located. Due to practical difficulties associated with measuring or estimating $C_c$, the partial pressure of $CO_2$ in the intercellular spaces ($C_i$) is often used instead, especially in the context of analyzing experimentally measured $CO_2$ response curves. Setting $C_c = C_i$ is equivalent to assuming infinite mesophyll conductance. Thus, the distinction between $C_i$ and $C_c$ is related to resistances along the path taken by $CO_2$ as it diffuses to Rubisco. It has no bearing on the response of Rubisco to $CO_2$, and hence no impact on the essential features of the min-*W* or min-*A* variants.
- Some sources express gas concentrations as mole fractions rather than partial pressures. This will change the numeric values of several parameters, but will have no impact on the general model behavior.
- In some sources, the non-photorespiratory $CO_2$ release term is not included in Equations **2a-c**; in this case, Equation **2d** becomes $A_n = \min\{A_c{}', A_j{}', A_p{}'\} - R_L$ or $A_n{}' = \min\{A_c{}', A_j{}', A_p{}'\}$, where $A_n{}' = A_n + R_L$ (and likewise for $A_c{}'$, $A_j{}'$, and $A_p{}'$). These versions all produce identical results to our Equation **2**.
- Several different versions of Equation **1b** (the equation for $W_j$) can be found in the literature, where the only changes are to the coefficients in the denominator (4 and 8 in our version). These coefficients are related to energy requirements involved with the electron transport chain, and different estimates exist for these requirements (Farquhar and von Caemmerer 1982; von Caemmerer 2000). Because of these differences, the coefficients can sometimes be considered as variables in the equation instead of taking fixed values.
- A simple minimum (Equation **1d**) is the most common way to choose between potential rates, but it produces sudden transitions between the different limitations. Sometimes alternatives to a simple minimum are used, such as quadratic mixing methods that produce smoother transitions (Kirschbaum and Farquhar 1984; Duursma 2015). Differences in $A_n$ between the variants will persist when using quadratic mixing.
- A more complex version of the FvCB model separately includes glycolate carbon leaving the photorespiratory pathway as glycine, serine, and 5,10-methylene-tetrahydrofolate.

This updated version does not affect our major conclusions, apart from new terms added to some equations and $C$ thresholds (Supplemental Section **S1.2**).

- Some sources describing 1-$\alpha$ versions of the model refer to the parameter as $\alpha$. Here we use the $\alpha_{old}$ notation introduced in Busch, Sage, and Farquhar (2018), which is a change in notation but the meaning of the parameter.
- Some sources implicitly assume $\alpha_{old} = 0$ when calculating TPU limited carboxylation or assimilation rates. This does not affect the conclusion that TPU cannot be limiting for $C \leq \Gamma^* \cdot (1 + 3 \cdot \alpha_{old})$, but the threshold simplifies to $C \leq \Gamma^*$ in this case.
- Instead of considering $W_p$ to be infinite for $C \leq \Gamma^* \cdot (1 + 3 \cdot \alpha_{old})$, some sources use $V_c = \min\{W_c, W_j\}$ for $C \leq \Gamma^* \cdot (1 + 3 \cdot \alpha_{old})$, only allowing $V_c = W_p$ for larger $C$. This ultimately produces the same effect of preventing TPU limitations in the forbidden range.
- The Rubisco limited carboxylation rate $W_c$ is sometimes referred to as the RuBP saturated carboxylation rate. Similar terminology is also sometimes used for $A_c$.
- Some sources do not consider TPU limitations at all, but the differences between the min-*W* and min-*A* approaches persist even without TPU.
- Some sources refer to $A_j$ as the electron transport limited (or light limited) assimilation rate, rather than the RuBP regeneration limited assimilation rate (and likewise for the associated carboxylation rate, $W_j$).
- Some sources refer to the rate of day respiration or mitochondrial respiration ($R_d$) rather than the rate of non-photorespiratory $CO_2$ release in the light ($R_L$). These quantities are numerically identical, but the $R_L$ notation is preferred to avoid any confusion with the rate of dark respiration, which is also often denoted by $R_d$. Furthermore, it is now understood that not all of this $CO_2$ release is due to mitochondrial respiration (Xu et al. 2021).

#### S1.2 The 3-α FvCB Model

A more complex version of the FvCB model separately includes glycolate carbon leaving the photorespiratory pathway as glycine, serine, and 5,10-methylene-tetrahydrofolate ($CH_2$-THF). (Busch, Sage, and Farquhar 2018; Busch 2020). This version of the model uses three different $\alpha$ parameters, each corresponding to one of these compounds: $\alpha_G$, $\alpha_S$, and $\alpha_T$, respectively.

Accounting for these separate sinks of glycolate carbon does not alter the Rubisco-limited carboxylation rate (Equation **1a**), but it does alter the energy requirements for RuBP regeneration (changing Equation **1b**) and the stoichiometry of inorganic phosphate (changing Equation **1c**). In the new model, these rates become:

$$W_c = \frac{C \cdot V_{c,max}}{C + K_C \cdot \left(1 + \frac{O}{K_O}\right)} \quad \textbf{(S1a)}$$

$$W_j = \frac{C \cdot J}{4 \cdot C + 8 \cdot \Gamma_{GT}^{*} (1 + 2 \cdot \alpha_G - \alpha_T + \alpha_S)} \quad \textbf{(S1b)}$$

$$W_p = \begin{cases} \infty, & C \le \Gamma_{GT}^{*} \cdot (1 + 3 \cdot \alpha_G + 6 \cdot \alpha_T + 4 \cdot \alpha_S) \\ \frac{3 \cdot C \cdot T_p}{C - \Gamma_{GT}^{*} \cdot (1 + 3 \cdot \alpha_G + 6 \cdot \alpha_T + 4 \cdot \alpha_S)}, & C > \Gamma_{GT}^{*} \cdot (1 + 3 \cdot \alpha_G + 6 \cdot \alpha_T + 4 \cdot \alpha_S) \end{cases} \quad \textbf{(S1c)}$$

where $\Gamma_{GT}^{*} = \Gamma^{*} \cdot (1 - \alpha_G + 2 \cdot \alpha_T)$ is the effective value of $\Gamma^{*}$. Glycolate carbon remaining in the cytosol as glycine or $CH_2$-THF alters the amount of $CO_2$ released during photorespiration, requiring the replacement of $\Gamma^{*}$ by $\Gamma_{GT}^{*}$, which also alters Equation **1e**. Thus, the new version of the model also includes

$$V_c = \min\{W_c, W_j, W_p\} \quad \textbf{(S1d)}$$

$$A_n = V_c \cdot \left(1 - \frac{\Gamma_{GT}^{*}}{C}\right) - R_L \quad \textbf{(S1e)}$$

Although the $C$ threshold in Equation **S1c** is not discussed in the original publications describing this model, it can nevertheless be derived following the same reasoning used in Section **3.3** of the main text. Fully written out, the threshold is $\Gamma^{*} \cdot (1 - \alpha_G + 2 \cdot \alpha_T) \cdot (1 + 3 \cdot \alpha_G + 6 \cdot \alpha_T + 4 \cdot \alpha_S)$.

Following the same reasoning used in Section **2.2** of the main text, it is possible to define a min-*A* version of Equation **S1**, which can also be considered as a 3-$\alpha$ version of Equation **2** of the main text:

$$A_c = W_c \cdot \left(1 - \frac{\Gamma_{GT}^{*}}{C}\right) - R_L = \frac{(C - \Gamma_{GT}^{*}) \cdot V_{c,max}}{C + K_C \cdot \left(1 + \frac{O}{K_O}\right)} - R_L. \quad \textbf{(S2a)}$$

$$A_j = W_j \cdot \left(1 - \frac{\Gamma_{GT}^{*}}{C}\right) - R_L = \frac{(C - \Gamma_{GT}^{*}) \cdot J}{4 \cdot C + 8 \cdot \Gamma_{GT}^{*} (1 + 2 \cdot \alpha_G - \alpha_T + \alpha_S)} - R_L, \quad \textbf{(S2b)}$$

$$A_p = W_p \cdot \left(1 - \frac{\Gamma^*_{GT}}{C}\right) - R_L = \frac{3 \cdot \left(C - \Gamma^*_{GT}\right) \cdot T_p}{C - \Gamma^*_{GT} \cdot (1 + 3 \cdot \alpha_G + 6 \cdot \alpha_T + 4 \cdot \alpha_S)} - R_L. \qquad (\mathbf{S2c})$$

$$A_n = \min\{A_c, A_j, A_p\}. \qquad (\mathbf{S2d})$$

Following similar reasoning as in Section **3.1** of the main text, it can be shown that Equations **S1** and **S2** disagree when $C < \Gamma^*_{GT} = \Gamma^* \cdot (1 - \alpha_G + 2 \cdot \alpha_T)$.

Thus, although Equations **S1** and **S2** represent a more complete understanding of photorespiration than Equations **1** and **2** of the main text, they also show that the 3-$\alpha$ version of the FvCB model is still subject to differences between the min-*W* and min-*A* approaches, and it still exhibits a biochemically-derived threshold for TPU limitations. In the main text, we use the 1-$\alpha$ version for clarity, because the equations are simpler.

### S2. Literature Survey

#### S2.1 Origins of the Min-*A* Approach

It is difficult to say with certainty when the min-*A* approach first appeared in the literature. The earliest instance of choosing a minimal assimilation rate instead of a minimal carboxylation rate in the context of the FvCB model known to the authors is Collatz et al. (1990). Equation 3 in that paper reads $J_c = \min\{W_e, W_c\}$, where $J_c$ represents the photosynthetic contribution to the net $CO_2$ assimilation rate (in other words, $A_n = J_c - R_L$). In this notation, $W_e$ and $W_c$ are therefore similar to $A_j + R_L$ and $A_c + R_L$ in our notation, respectively.

Another early instance, which more closely aligns with the min-*A* approach as presented here, can be found in Collatz et al. (1991). Equation 2 in that paper reads $A_n \approx \min\{J_E, J_C, J_S\} - R_d$, where $J_E$, $J_C$, and $J_S$ are the photosynthetic contributions to the net $CO_2$ assimilation rate when carboxylation is limited by light, Rubisco, or sucrose synthesis; these rates are similar to $A_j + R_L$, $A_c + R_L$, and $A_p + R_L$ in our notation, respectively. Although these equations are presented as being equivalent to those in Kirschbaum and Farquhar (1984), there are several key differences (Walker et al. 2021).

#### S2.2 Highly-Cited Papers Published 1990 – 2023

To make an estimate of the relative prevalence of FvCB model variants in peer-reviewed scientific papers, we used the Web of Science Core Collection (accessed through the University of Illinois library between March 21, 2023 and April 7, 2023) to identify all documents within the database that cite the 1980 FvCB paper and were published in 1990 or later; 5672 records met these criteria. (This range of publication dates was chosen to exclude the period before the min-*A* variant was present in the literature.) These papers were sorted according to their own citations. Starting with the highest-cited paper, each one was manually inspected to identify publications that include at least one FvCB model equation and clearly indicate a method for choosing a rate-limiting process. Papers were investigated this way until fifteen had been identified; then these papers were categorized according to whether they presented the min-*W* approach, the min-*A* approach, or another variant (Table **S1**). One popular textbook was also included; its citation count was determined using Google Scholar since it was not present in the Web of Science database. The results from this analysis are summarized in Figure **1** of the main text.

Many of the papers identified this way include Equation **1d** or **2d** (as defined in the main text), making categorization straightforward. However, several of the papers require additional explanation to justify their categorization:

*von Caemmerer (2000)*: In this textbook, the author writes, "$A$ proceeds at a minimum of the three rates $A_c$, $A_j$, $A_p$. This is illustrated in Fig. 2.6." Nevertheless, a close look at Figure 2.6 shows that the net assimilation rate is set by $A_c$ at low $CO_2$ partial pressures rather than $A_j$, even though $A_j$ is smaller. In fact, an attempt to understand the discrepancy between Equation

2.27 and Figure 2.6 in this textbook was the original impetus for initiating the analysis that led to this manuscript.

*Sharkey et al. (2007):* In this paper, the authors write, "Equations 1, 2 and 3 can be put into a spread sheet and $\mathrm{V_{c,max}}$, $\mathrm{J}$ and $\mathrm{TPU}$ can be adjusted manually until each modelled line meets or exceeds all of the data points." Although this is not an equation, it clearly expresses the idea that the experimental measurements should be compared with the minimum of $A_c$, $A_j$, and $A_p$, which places this paper in the min-*A* category.

*Stitt (1991):* In this paper, the author writes, "Although it has sometimes been assumed or asserted that photosynthesis is controlled by single "limiting" factors… this assertion is not easy to reconcile with the highly interactive and complex regulation of photosynthesis… nor with the experimental evidence that control is shared between several enzymes and redistributes in a very flexible manner in other pathways." This skepticism about whether the processes considered in the FvCB model are sufficient to explain the regulation of photosynthesis prevents it from being classified as a pure expression of either the min-*W* or min-*A* approach.

*de Pury and Farquhar (1997):* In this paper, the expressions for Rubisco and electron transport limited assimilation (Equations 2 and 4 in the paper) are restricted to $C > \Gamma^*$, which is exactly the range where the variants agree. So, although Equation 1 in the paper chooses a minimal assimilation rate, this paper cannot be classified as expressing either variant.

*Harley et al. (1992):* In this paper, the authors write, "Current models of $C_3$ leaf photosynthesis assume that carboxylation of RuBP by Rubisco is limited by one of three factors... These are (a) the activity of Rubisco, (b) the regeneration of RuBP, or (c) the release of phosphate during the metabolism of triose phosphate to either starch or sucrose." Although this is not an equation, it clearly expresses the idea of choosing a limiting carboxylation rate, which places this paper in the min-*W* category.

| Publication | Citations | Variant Presented | Justification |
|---|---|---|---|
| Collatz et al. (1991) | 1602 | Min-*A* | Equation 2 |
| Krinner et al. (2005) | 1403 | Min-*W* | Equations A2 and A3 |
| Sellers et al. (1996) | 1358 | Min-*A* | Equation 17 |
| von Caemmerer (2000) | 1318* | Min-*A* | Equation 2.27 |
| Leuning (1995) | 1127 | Min-*A* | Equation 10 |
| Sharkey et al. (2007) | 903 | Min-*A* | *(see above)* |
| Foley et al. (1996) | 901 | Min-*A* | Equation 1 |
| Stitt (1991) | 859 | Other | *(see above)* |
| Pury and Farquhar (1997) | 856 | Other | *(see above)* |
| Long and Bernacchi (2003) | 832 | Min-*W* | Equation 6 |
| Wullschleger (1993) | 825 | Min-*W* | Equation 2 |
| Long (1991) | 823 | Min-*W* | Equation 1 |
| Sellers et al. (1992) | 712 | Min-*A* | Equations 14a and 14b reduce to a simple minimum when $\theta = \beta = 1$ |
| Haxeltine and Prentice (1996) | 711 | Min-*A* | Equation 2 reduces to a simple minimum when $\theta = 1$ |
| Harley et al. (1992) | 686 | Min-*W* | *(see above)* |
| Medlyn et al. (2002) | 588 | Min-*A* | Equation 1 |

*Table S1: Highly-cited papers published in 1990 or later that express one or more FvCB model equations and clearly indicate a method for choosing a rate-limiting process. Citation counts were obtained from the Web of Science database or Google Scholar (marked by an asterisk (*)). References to equations in the justification column use each paper's own numbering scheme.*

#### S2.3 Highly-Cited Papers Published 2013 – 2023

The most recent paper identified in the previous section was published in 2007, so that collection may not be representative of newer papers. To address this, we repeated the same procedure for papers published in the last ten years, identifying the top six papers (Table **S2**). The results from this analysis are summarized in Figure **1** of the main text. As before, many of the papers can be categorized based on one or two equations, but one requires additional explanation:

*Franks et al. (2014):* In this paper, the authors write, "Briefly, in the Farquhar-von Caemmerer-Berry biochemical model for photosynthesis... $\mathrm{A_n} = \min[\mathrm{W_e}, \mathrm{W_c}]$ where $\mathrm{W_e}$ is the light-limited rate and $\mathrm{W_c}$ is the Rubisco capacity-limited rate." Although this is not a numbered equation in the paper, it clearly expresses the net $CO_2$ assimilation rate as the minimum of two potential net $CO_2$ assimilation rates, which places this paper in the min-*A* category.

| Publication | Citations | Variant Presented | Justification |
|---|---|---|---|
| Yamori, Hikosaka, and Way (2014) | 570 | Min-*W* | Equation 3 in Supplementary Material 1 |
| Smith and Dukes (2013) | 306 | Min-*A* | Equation 1 |
| Walker et al. (2014) | 256 | Min-*W* | Equation 2 |
| Duursma (2015) | 248 | Min-*A* | Equation 1 |
| Bonan et al. (2014) | 217 | Min-*A* | Equation A10 |
| Franks et al. (2014) | 156 | Min-*A* | *(see above)* |

*Table S2: Highly-cited papers published in 2013 or later that express one or more FvCB model equations and clearly indicate a method for choosing a rate-limiting process. Citation counts were obtained from the Web of Science database. References to equations in the justification column use each paper's own numbering scheme.*

#### S2.4 Tools for Estimating FvCB Model Parameters from $CO_2$ Response Curves

Besides a general survey of impactful papers, we also categorized a set of tools and procedures for estimating FvCB model parameters from experimentally measured $CO_2$ response curves (Table **S3**). Most of these tools and procedures come from peer-reviewed publications; the only exception is the Excel spreadsheet from *landflux.org*, which was included because it has been used in several publications, such as Croft et al. (2017). Some publications describe more than one tool; in this case, each one is counted separately. Many strategies have been developed for parameter estimation from $CO_2$ response curves:

- *Manual*: In manual fitting methods, the user directly specifies the rate-limiting process (either Rubisco activity, RuBP regeneration, or TPU) for each point in a $CO_2$ response curve. This is distinct from either the min-$W$ or min-$A$ variants, because it essentially excludes Equations **1e** and **2d** of the main text; however, users typically specify sequences of limitations that are possible in the min-$W$ variant rather than the min-$A$ variant.
- *Exhaustive*: Exhaustive methods are similar to manual methods in the sense that a rate-limiting process is assigned to each point in a $CO_2$ response curve before any fitting occurs. The difference is that an algorithm is used to test all possible ways to assign the rate-limiting processes, and the choice with the closest fit is chosen as being optimal. Again, this is distinct from the min-$W$ and min-$A$ variants, although the sequences of limitations are typically limited to (a subset of) those possible in the min-$W$ variant.
- *Whole-curve*: Whole-curve methods determine the rate-limiting process at each point in a $CO_2$ response curve by choosing either a minimal carboxylation rate or a minimal net assimilation rate. Tools and procedures using whole-curve methods can be categorized as described in the main text (min-$W$ variant, min-$A$ variant, min-$A$+FRL variant, etc.).

The results from this analysis are summarized in Figure **1** of the main text. Some of the entries in this section require additional explanation:

*Duursma (2015):* The plantecophys R package actually provides three different fitting methods, which can be specified using the `fitmethod` argument of the `fitaci` function. Values of `default`, `bilinear`, and `onepoint` correspond to min-$A$+FRL+FTT, exhaustive, and manual methods, respectively. Note that the publication only discusses a whole-curve fitting method based on the min-$A$ approach; although the `default` method from the package is similar, it is not exactly the same due to the presence of code modifications that actually implement the min-$A$+FRL+FTT variant.

*Gu et al. (2010):* This fitting method is publically available, but only through a web service at https://leafweb.org/. It is not possible to access the source code, so we assume it follows the exhaustive method described in the associated publication. This paper expresses the min-$W$ variant equations—including the limited range of $C$ where TPU limitations are possible—and discusses the possible sequences of rate-limiting processes in the model. It also includes a short discussion of the min-*A* variant (without using that name). However, these issues are not fully explored, the FRL and FTT modifications are not mentioned, and the min-*A* variant with or without the FRL and FTT modifications are not explicitly connected to possible errors in parameter estimation.

*Parsons et al. (1997):* A compiled executable program associated with this publication is available online via the Internet Archive; however, it will not run on modern computer systems. Since it is not possible to access the source code or run the program, we simply assume it uses the min-*W* equations as described in the associated publication.

*Moualeu-Ngangue, Chen, and Stützel (2017):* The SAS code provided with this publication uses the min-*A* variant for fitting but the min-*A*+FRL variant for displaying the results of the fit.

*Stinziano et al. (2021):* Although fitting is performed using an exhaustive approach, the min-*A*+FRL variant is used when displaying the resulting fits.

*Xiao et al. (2021):* The R code used in the Jupyter notebook chooses a minimal assimilation rate for $C \geq \Gamma^*$ and a maximal assimilation rate for $C < \Gamma^*$, which is equivalent to the min-*W* equations as presented in the main text.

*Lochocki, Salesse-Smith, and McGrath (2025)*: The PhotoGEA R package was added to the list after the initial literature survey, since it was not published until 2025. It uses the min-*W* approach by default, but the min-*A* approach can be enabled through an optional input argument, as well as the FRL and FTT modifications; these options are for illustrative purposes only and are not intended to be used for most fits.

| Publication | Times Cited | Variant Presented | Code Source | Code version | Tool Type | Limiting Process Determination |
|---|---|---|---|---|---|---|
| Sharkey et al. (2007) | 903 | Min-*A* | SI | 2007.1 | Excel spreadsheet | Manual |
| Long and Bernacchi (2003) | 831 | Min-*W* | -- | -- | Written procedure | Manual |
| Ethier and Livingston (2004) | 448 | Min-*A* | -- | -- | Written procedure | Min-*A* |
| Duursma (2015) | 248 | Min-*A* | *(a)* | 1.4-6 | R package | Min-*A*+FRL+FTT |
| | | | | | | Exhaustive |
| | | | | | | Manual |
| Gu et al. (2010) | 130 | Min-*W* | NA | -- | Web service | Exhaustive* |
| Sharkey (2016) | 110 | -- | SI | 2.0 | Excel spreadsheet | Manual |
| Dubois et al. (2007) | 104 | Min-*A* | SI | -- | SAS code | Min-*A* |
| Bellasio et al. (2016) | 51 | -- | SI | EFT33_2015 | Excel spreadsheet | Manual |
| Parsons et al. (1997) | 43 | Min-*W* | *(b)* | -- | Spreadsheet | Min-*W** |
| Moualeu-Ngangue, Chen, and Stützel (2017) | 33 | Min-*A* | SI | 21 October 2016 | Excel spreadsheet | Min-*A* |
| | | | | | SAS code | Min-*A* |
| Gregory et al. (2021) | 7 | -- | *(c)* | 1.2.0 | R package | Min-*W*+FTT |
| Stinziano et al. (2021) | 4 | -- | *(d)* | 2.1.2 | R package | Exhaustive |
| Xiao et al. (2021) | 3 | Min-*W* | *(e)* | 1 | Jupyter notebook | Min-*W* |
| -- | -- | -- | *(f)* | 10.1 | Excel spreadsheet | Min-*A* |
| Lochocki, Salesse-Smith, and McGrath (2025) | 0 | Min-*W* | *(g)* | 1.2.0 | R package | Min-*W* |

*Table S3: Publically-available tools or procedures for estimating FvCB model parameters from $CO_2$ response curves. The "Variant Presented" column refers to the equations as they appear in the publication associated with each tool. Asterisks (*) in the "Limiting Process Determination" column indicate presumptive guesses based on the equations presented in the associated publication when source code is not available. The entries in the "Code Source" column mean: SI = Supplementary information associated with the publication; NA = Not publically available; (a) https://github.com/RemkoDuursma/plantecophys; (b) https://web.archive.org/web/19970717042147/http://www.dundee.ac.uk/BioScience/photosyn.htm; (c) https://github.com/poales/msuRACiFit; (d) https://github.com/cdmuir/photosynthesis; (e) https://github.com/xiaoyizz78/FvCB-JAGS; (f) https://landflux.org/; (g) https://github.com/eloch216/PhotoGEA. Citation counts were obtained from the Web of Science database.*

#### S2.5 Lower C Threshold for TPU

Of the thirty-two distinct publications identified in the survey described in Sections **S2.2**-**S2.4** above, fourteen present an equation for TPU-limited carboxylation or assimilation (Table **S4**). Many of these include statements implying that TPU can only limit carboxylation or assimilation at higher values of $C$, such as “there is now increasing evidence that limitations imposed on the assimilation of $CO_2$ by triose phosphate utilization may be a common occurrence under conditions of high $C_i$ and high irradiance” (Wullschleger 1993). However, only two publications explicitly includes a lower $C$ threshold for TPU limitations in the equations themselves: Gu et al. (2010) and Lochocki, Salesse-Smith, and McGrath (2025).

| Publication | Variant Presented | Lower C Threshold for TPU |
|---|---|---|
| Collatz et al. (1991) | Min-*A* | -- |
| Sellers et al. (1992) | Min-*A* | -- |
| Wullschleger (1993) | Min-*W* | -- |
| Foley et al. (1996) | Min-*A* | -- |
| Sellers et al. (1996) | Min-*A* | -- |
| Parsons et al. (1997) | Min-*A* | -- |
| von Caemmerer (2000) | Min-*A* | -- |
| Long and Bernacchi (2003) | Min-*W* | -- |
| Dubois et al. (2007) | Min-*A* | -- |
| Sharkey et al. (2007) | Min-*A* | -- |
| Gu et al. (2010) | Min-*W* | $\Gamma^* \cdot (1 + 3 \cdot \alpha_{old})$ |
| Bellasio, Beerling, and Griffiths (2016) | -- | -- |
| Moualeu-Ngangue, Chen, and Stützel (2017) | Min-*A* | -- |
| Lochocki, Salesse-Smith, and McGrath (2025) | Min-*W* | $\Gamma^* \cdot (1 + 3 \cdot \alpha_{old})$ or $\Gamma_{GT}^* \cdot (1 + 3\alpha_G + 6\alpha_T + 4\alpha_S)$ |

*Table S4: Publications from Tables S1-S3 that include equations for TPU-limited assimilation or carboxylation. The “Variant Presented” and “Lower C Threshold for TPU” columns refer to the equations as they appear in the publication, and do not reflect any equations implemented in computer code associated with the publications.*

### S3. $CO_2$ Response in the Min-*W* Variant

#### S3.1. Handling C = 0 in the Min-*W* Variant

Equation **1e** cannot be evaluated at $C = 0$ because doing so would require a division by zero (due to the $\Gamma^*/C$ term). However, Equations **2a** and **2b** can be evaluated at $C = 0$; in this case, they become

$$A_c|_{C=0} = -\frac{\Gamma^* \cdot V_{c,max}}{K_C \cdot \left(1 + \frac{O}{K_O}\right)} - R_L \tag{A1}$$

and

$$A_j|_{C=0} = -\frac{J}{8} - R_L. \tag{A2}$$

Then, the net $CO_2$ assimilation rate at $C = 0$ can be determined using Equation **3**:

$$A_n|_{C=0} = \max\{A_c|_{C=0}, A_j|_{C=0}\}. \tag{A3}$$

Note that there is no need to consider TPU limitations at $C = 0$ (Section **3.3**).

The expressions for $A_c|_{C=0}$ and $A_j|_{C=0}$ from Equations **A1** and **A2** can be equated to find the value of $J$ where the two rates are equal ($J_{eq}$):

$$J_{eq} = \frac{8 \cdot \Gamma^* \cdot V_{c,max}}{K_C \cdot \left(1 + \frac{O}{K_O}\right)}. \tag{A4}$$

If $J > J_{eq}$, then $A_j|_{C=0} < A_c|_{C=0}$; in this case, carboxylation is Rubisco limited at low $C$. Likewise, carboxylation is RuBP regeneration limited at low $C$ when $J < J_{eq}$.

#### S3.2. The Crossover Between Rubisco and RuBP Regeneration Limitations in the Min-*W* Variant

The crossover value of $C$ where $W_c$ and $W_j$ are equal ($C_{cj}$) can be found by equating Equations **1a** and **1b** and solving for the $CO_2$ partial pressure:

$$C_{cj} = \frac{K_C \cdot \left(1 + \frac{O}{K_O}\right) \cdot \frac{J}{4 \cdot V_{c,max}} - \frac{8}{4} \cdot \Gamma^*}{1 - \frac{J}{4 \cdot V_{c,max}}}. \tag{A5}$$

Note that even though this equation contains terms of $J/(4 \cdot V_{c,max})$, it can be evaluated at $V_{c,max} = 0$. In this case it reduces to $C_{cj} = -K_c \cdot (1 + O/K_o)$, which is always negative. Since $C$ cannot be negative (by definition), this simply indicates that a crossover does not occur when $V_{c,max} = 0$.

Also note that when $\frac{J}{V_{c,max}} = \frac{(8+4) \cdot \Gamma^*}{K_C \cdot \left(1 + \frac{O}{K_O}\right) + \Gamma^*}$, Equation **A5** evaluates to $C_{cj} = \Gamma^*$. In other words, there is a special case where the crossover happens to occur exactly at $\mathrm{C} = \Gamma^*$ in the min-*W* variant. This situation only occurs at a very precise ratio of $\frac{J}{V_{c,max}}$ (approximately 0.7701 using the parameter values in Table **1**), so it is unlikely to be observed in practice.

In general, the rational function defined in Equation **A5** can produce negative values of $C_{cj}$ whenever the numerator and denominator have opposite signs. The numerator is zero when $J$ is given by

$$J_n = \frac{8 \cdot \Gamma^* \cdot V_{c,max}}{K_C \cdot \left(1 + \frac{O}{K_O}\right)}, \quad \text{(A6)}$$

while the denominator is zero when $J$ is given by

$$J_d = 4 \cdot V_{c,max}. \quad \text{(A7)}$$

For fixed values of $\Gamma^*$, $V_{c,max}$, $K_C$, $K_O$, and $O$, the two curves defined by Equations **A6** and **A7** therefore form the boundaries of the regions of $(V_{c,max}, J)$-space where a crossover is not possible due to negative $C_{cj}$.

The numerator is negative whenever $J < J_n$ and the denominator is negative whenever $J > J_d$. It is possible for both conditions to be satisfied, but this would require $\Gamma^* > 4/8 \cdot k_c \cdot (1 + O/K_o)$, a situation that is unlikely to occur in most situations because $k_c$ is usually much larger than $\Gamma^*$. So, in most cases, negative values of $C_{cj}$ occur for $J > J_n$ or $J < J_d$. Since $J_n$ is larger than $J_d$, there is a region of $J$ values between $J_n$ and $J_d$ where a crossover is possible. Note that the equations for $J_n$ (Equation **A6**) and $J_{eq}$ (Equation **A4**) are identical so that $J_n \equiv J_{eq}$; from this, we can see that when carboxylation is RuBP regeneration limited at $C = 0$ with typical parameter values, a crossover to Rubisco limited carboxylation never occurs; all crossovers must be from Rubisco limited to RuBP regeneration limited carboxylation. Nevertheless, other crossovers are mathematically possible with atypical parameter values (Supplemental Section **S4**).

The considerations above describe situations where no crossover occurs because it is mathematically impossible; however, there are other situations where no crossover occurs because it is impossible to observe in practice. For example, if $C_{cj}$ exceeds 2000 µbar, it is unlikely that such a crossover would ever be observed in a measured $CO_2$ response curve.

##### S3.3. An Example

Using the parameter values specified in Table **1**, we can determine the following:

- At $C = 0$, $A_c$ is given by $A_c|_{C=0}$ = -6.86 µmol m$^{-2}$ s$^{-1}$ and $A_j$ is given by $A_j|_{C=0}$ = -21.25 µmol m$^{-2}$ s$^{-1}$, so carboxylation is limited by Rubisco activity at low values of $C$.
- $W_c$ and $W_j$ become equal at $C_{cj}$ = 281.76 µbar, so there is a crossover from Rubisco limited to RuBP regeneration limited carboxylation at a reasonable $CO_2$ partial pressure.

Thus, we can summarize the sequence of rate-limiting processes corresponding to those values of $V_{c,max}$ and $J$ as $A_c \rightarrow A_j$.

#### S4. Sequences of Limiting Processes for High Γ* and/or Low Kc

For the parameter values in Table **1** in the main text, it was shown that only certain sequences of limiting processes are predicted by the min-*W* variant as $C$ increases (Figure **2**):

- $A_c$
- $A_c \rightarrow A_j$
- $A_c \rightarrow A_j \rightarrow A_p$
- $A_c \rightarrow A_p$
- $A_j$
- $A_j \rightarrow A_p$

As mentioned in Supplemental Section **S3**, other transitions are mathematically possible. These can happen when $\Gamma^* > 4/8 \cdot K_c \cdot (1 + O/K_o)$, which could occur in principle for very large $\Gamma^*$ or very small $K_c$. In this case, $J_d$ is larger than $J_n$, and negative values of $C_{cj}$ occur for $J < J_n$ or $J > J_d$. Because $J_n$ and $J_{eq}$ are identical, a crossover to RuBP regeneration limited carboxylation never occurs in this scenario; all crossovers must be from RuBP regeneration limited to Rubisco limited carboxylation. It may be the case that biochemical constraints prevent $\Gamma^* > 4/8 \cdot K_c \cdot (1 + O/K_o)$ from ever occurring in a real chloroplast, but the min-*W* variant does not expressly forbid it, and does allow for a crossover from $W_j$ to $W_c$.

Indeed, it is possible to choose parameters that satisfy this condition and then generate maps of the limiting processes as in Figure **2**. Making such maps, we find the appearance of two new transitions (Figure **S1**):

- $A_j \rightarrow A_c$
- $A_j \rightarrow A_c \rightarrow A_p$

Although these transitions do not seem to occur in nature, there is nothing in the FvCB model to exclude them. Thus, their absence in experimentally measured $CO_2$ response curves is due to restrictions on Rubisco properties and environmental conditions rather than something inherent to the processes of Rubisco limited or RuBP regeneration limited carboxylation.

Note that the transitions listed in this section cover all possible transitions in the min-*W* variant. Because $W_c$ and $W_j$ are monotonic increasing functions of $C$ and $W_p$ is a monotonic decreasing function of $C$, it is not possible for Rubisco or RuBP regeneration to become limiting at higher $CO_2$ concentrations once TPU becomes limiting. In other words, sequences like $A_c \rightarrow A_p \rightarrow A_j$ are forbidden. Furthermore, Equation **A6** has just one solution for fixed values of the photosynthetic parameters, so there can be at most one crossover between $A_c$ and $A_j$. In other words, sequences like $A_c \rightarrow A_j \rightarrow A_c$ are forbidden. (Note that these forbidden sequences do occur in the min-*A* variant.)

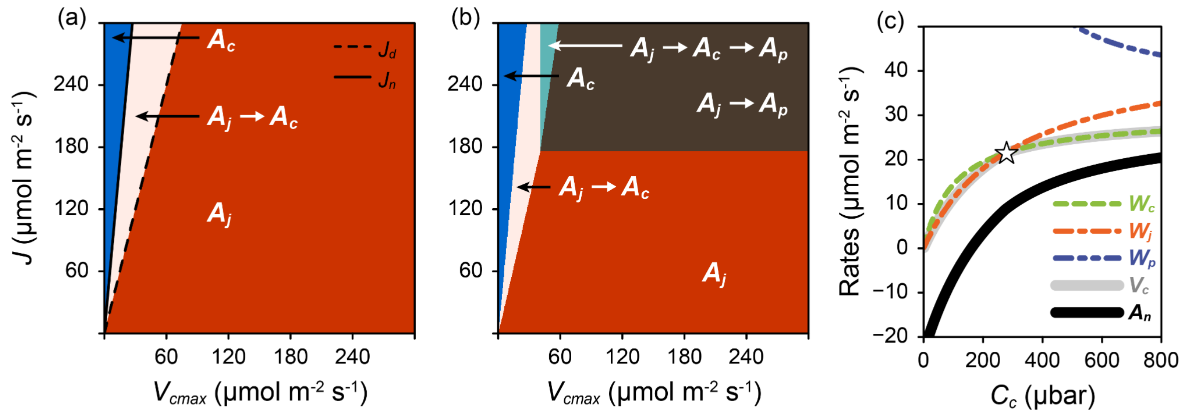

*Figure S1: Maps of $(V_{c,max}, J)$-space showing which sequences of limiting processes are predicted by the min-W variant as $C_c$ increases, either neglecting (a) or including (b) possible TPU limitations. (a) also includes $J_d$ and $J_n$ (Equations B2 and B3) as dashed and solid black lines, respectively. In (b), all labels refer to observed sequences below a threshold of 2000 µbar. (c) shows $W_c$, $W_j$, $W_p$, $V_c$ and $A_n$ as calculated by the min-W variant with $V_{c,max}$ = 100 µmol m$^{-2}$ s$^{-1}$ and $J$ = 10 µmol m$^{-2}$ s$^{-1}$. The crossover point where the rate-limiting process changes is marked with a white star. For these calculations, $\Gamma^*$ = 150 µbar and $K_c$ = 50 µbar; all other parameters used in (a-c) other than $V_{c,max}$ and $J$ were set to the values specified in Table 1.*

### S5. Fitting Experimentally Measured $CO_2$ Response Curves

#### S5.1 Measurement Details

*Nicotiana tabacum* cv. Samsun seeds were sown on January 17, 2023, and grown in the RIPE greenhouse at the University of Illinois Urbana-Champaign. $CO_2$ response curves were measured March 21 – 24, 2023, when the plants were 9 weeks old. These plants were grown during the Illinois winter, and hourly logs of incident photosynthetically active flux density (PPFD) indicate that daytime light levels most often fell within 200 – 350 µmol $m^{-2}$ $s^{-1}$ during the growth period. Almost half of all days during this period had maximum PPFD levels below 600 µmol $m^{-2}$ $s^{-1}$. The average daily minimum and maximum temperatures were 23.9 °C and 30.4 °C, respectively, and the median hourly temperature was 27.9 °C.

$CO_2$ response curves were obtained using Licor LI-6800 portable gas exchange measurement systems. Leaf temperature was set to 27 °C and the vapor pressure deficit in the leaf chamber was set to 1.3 kPa. To generate the response curves, the reference $CO_2$ concentration was varied in one of the following sequences: 400, 300, 200, 150, 100, 75, 50, 40, 30, 20, 10, 400, 400, 600, 800, 1000, 1200, 1500 ppm (on March 21) or 400, 300, 200, 150, 100, 75, 50, 40, 30, 20, 10, 400, 400, 500, 600, 800, 1000, 1200, 1500 (on March 23-24). Measurements were made using the youngest fully expanded leaf.

$CO_2$ response curves were measured at incident photosynthetically active photon flux density (PPFD) of 100, 150, 200, 250, 300, 400, 450, 500, 600, 800, 1000, 1200, or 1500 µmol $m^{-2}$ $s^{-1}$. Three or four curves were measured from different plants at each incident PPFD, with the exception of 450 and 1200 µmol $m^{-2}$ $s^{-1}$ (one curve each) and 1000 and 1500 µmol $m^{-2}$ $s^{-1}$ (two curves each), for a total of thirty-six curves.

These curves have been published previously (Lochocki, Salesse-Smith, and McGrath 2025), but the fitting approach used here differs from the original publication.

#### S5.2 All Curve Fits

The `fit_c3_aci` function from the *PhotoGEA* R package was used for all fits. By default, this function uses Equation **1**. To perform fits using Equation **6**, the optional `consider_depletion` input argument was set to TRUE. For fits using Equation **2** with the FTT modification, the optional `use_min_A` input argument was set to TRUE, and the optional `TPU_threshold` input argument was set to 400 µmol $mol^{-1}$.

Figure **5** in the main text includes curve fits at low $C_i$ for one curve with $A_d$ limitations (curve number 11, designated “600 - wt-9 - mcgrath1”) and one without $A_d$ limitations (curve number 29, designated “600 - wt-8 - mcgrath2”). In the following figures (Figures **S2** – **S9**), we show fits for all curves in the set, at low $C_i$ and across the entire measured range of $C_i$. The curve naming scheme include three parts: the incident PPFD (in µmol $m^{-2}$ $s^{-1}$, e.g. 600), the particular plant identifier (e.g. wt-9), and the machine used to measure the curve (e.g. mcgrath1). Curve names for each curve number are shown in Table **S5**.

| **Curves with $A_d$ limitations** | | **Curves without $A_d$ limitations** | |
|---|---|---|---|
| **Curve number** | **Curve identifier** | **Curve number** | **Curve identifier** |
| 1 | 1000 - wt-5 - mcgrath1 | 19 | 300 - wt-3 - ripe5 |
| 2 | 500 - wt-9 - mcgrath1 | 20 | 250 - wt-7 - mcgrath1 |
| 3 | 400 - wt-2 - mcgrath1 | 21 | 250 - wt-1 - ripe5 |
| 4 | 300 - wt-9 - ripe5 | 22 | 300 - wt-1 - mcgrath1 |
| 5 | 150 - wt-6 - ripe5 | 23 | 150 - wt-1 - ripe5 |
| 6 | 400 - wt-3 - ripe5 | 24 | 200 - wt-8 - mcgrath1 |
| 7 | 1000 - wt-1 - mcgrath1 | 25 | 450 - wt-5 - mcgrath2 |
| 8 | 400 - wt-9 - mcgrath1 | 26 | 200 - wt-1 - ripe5 |
| 9 | 250 - wt-5 - ripe5 | 27 | 150 - wt-3 - mcgrath1 |
| 10 | 100 - wt-7 - ripe5 | 28 | 400 - wt-10 - ripe5 |
| 11 | 600 - wt-9 - mcgrath1 | 29 | 600 - wt-8 - mcgrath2 |
| 12 | 1500 - wt-10 - mcgrath2 | 30 | 300 - wt-8 - ripe5 |
| 13 | 800 - wt-5 - mcgrath1 | 31 | 100 - wt-4 - ripe5 |
| 14 | 1500 - wt-7 - mcgrath1 | 32 | 200 - wt-10 - mcgrath1 |
| 15 | 500 - wt-3 - ripe5 | 33 | 500 - wt-10 - mcgrath1 |
| 16 | 1200 - wt-6 - mcgrath1 | 34 | 100 - wt-8 - mcgrath1 |
| 17 | 500 - wt-7 - ripe5 | 35 | 250 - wt-1 - mcgrath2 |
| 18 | 600 - wt-1 - mcgrath1 | 36 | 100 - wt-3-2 - mcgrath1 |

435 *Table S5: Curve numbers and identifiers for curves with and without $A_d$ limitations.*

#### S5.2.1 Min-*W* Variant (all $C_i$)

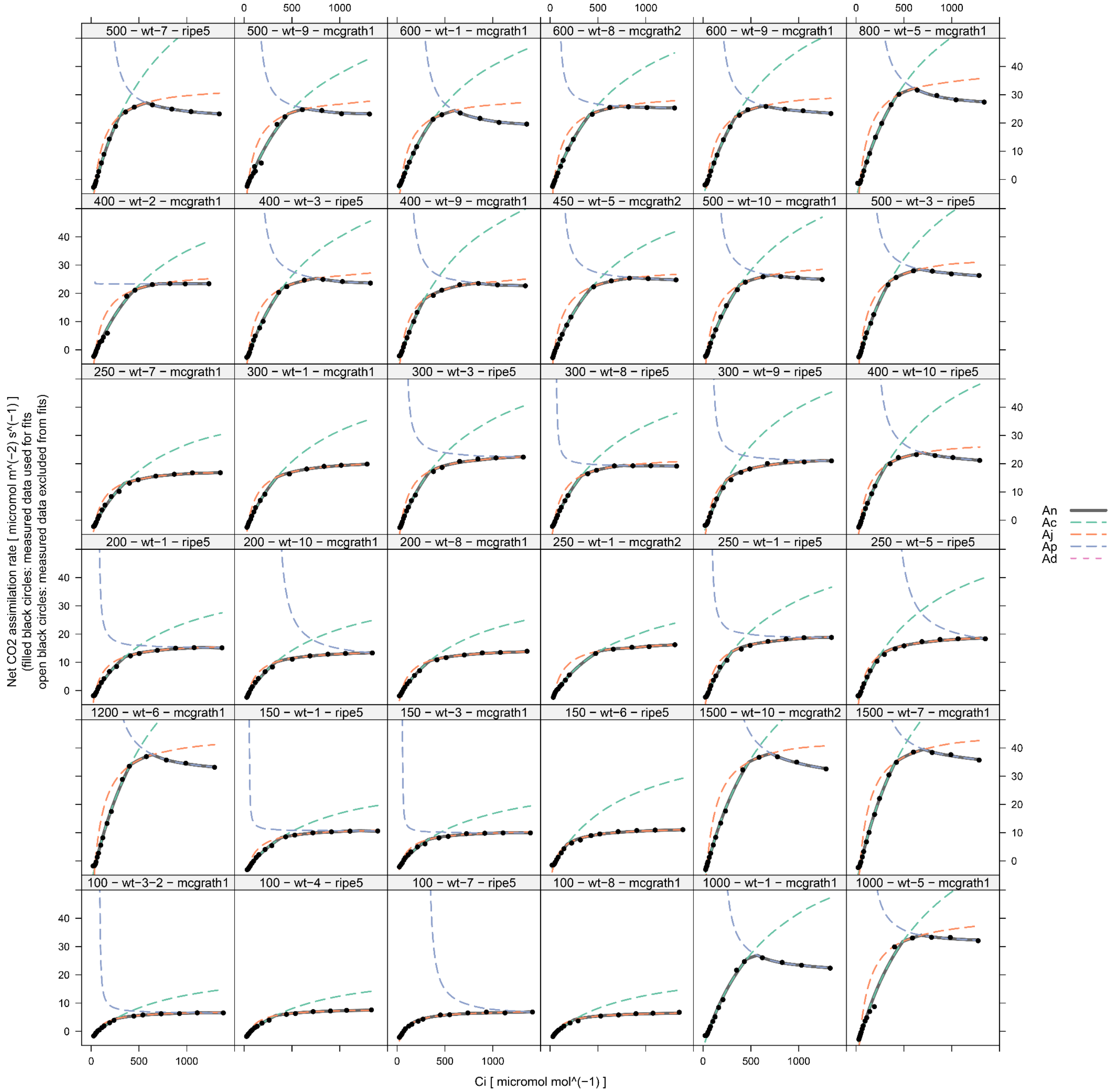

*Figure S2: Min-W variant (either Equation **1** or Equation **6**) curve fits made using all measured points, shown across the full range of $A_n$ and $C_i$.*

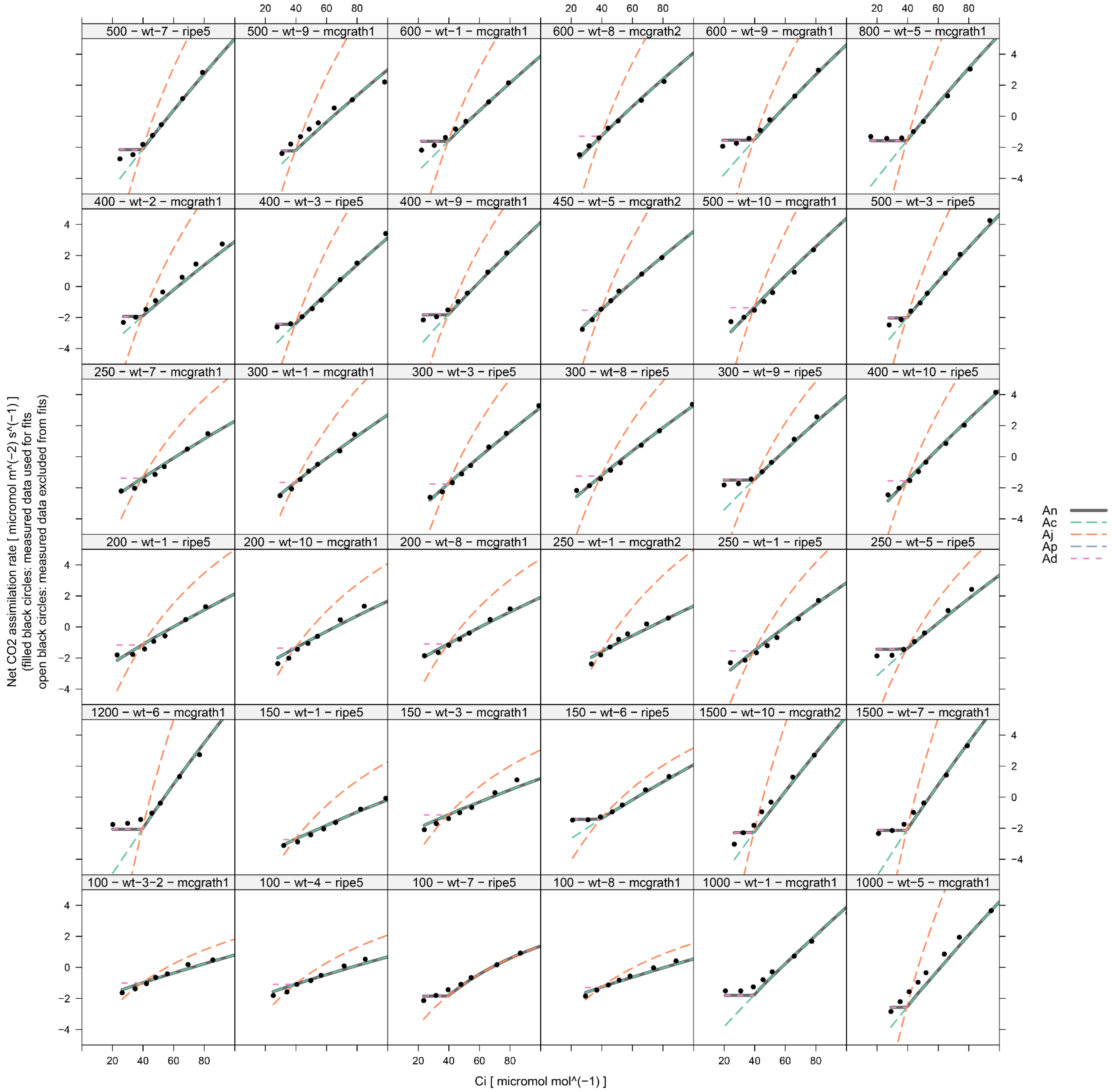

*Figure S3: Min-W variant (either Equation **1** or Equation **6**) curve fits made using all measured points, shown across a limited range of $A_n$ and $C_i$ in order to highlight the fits at low $C_i$.*

#### S5.2.1 Min-*A*+FTT variant (all $C_i$)

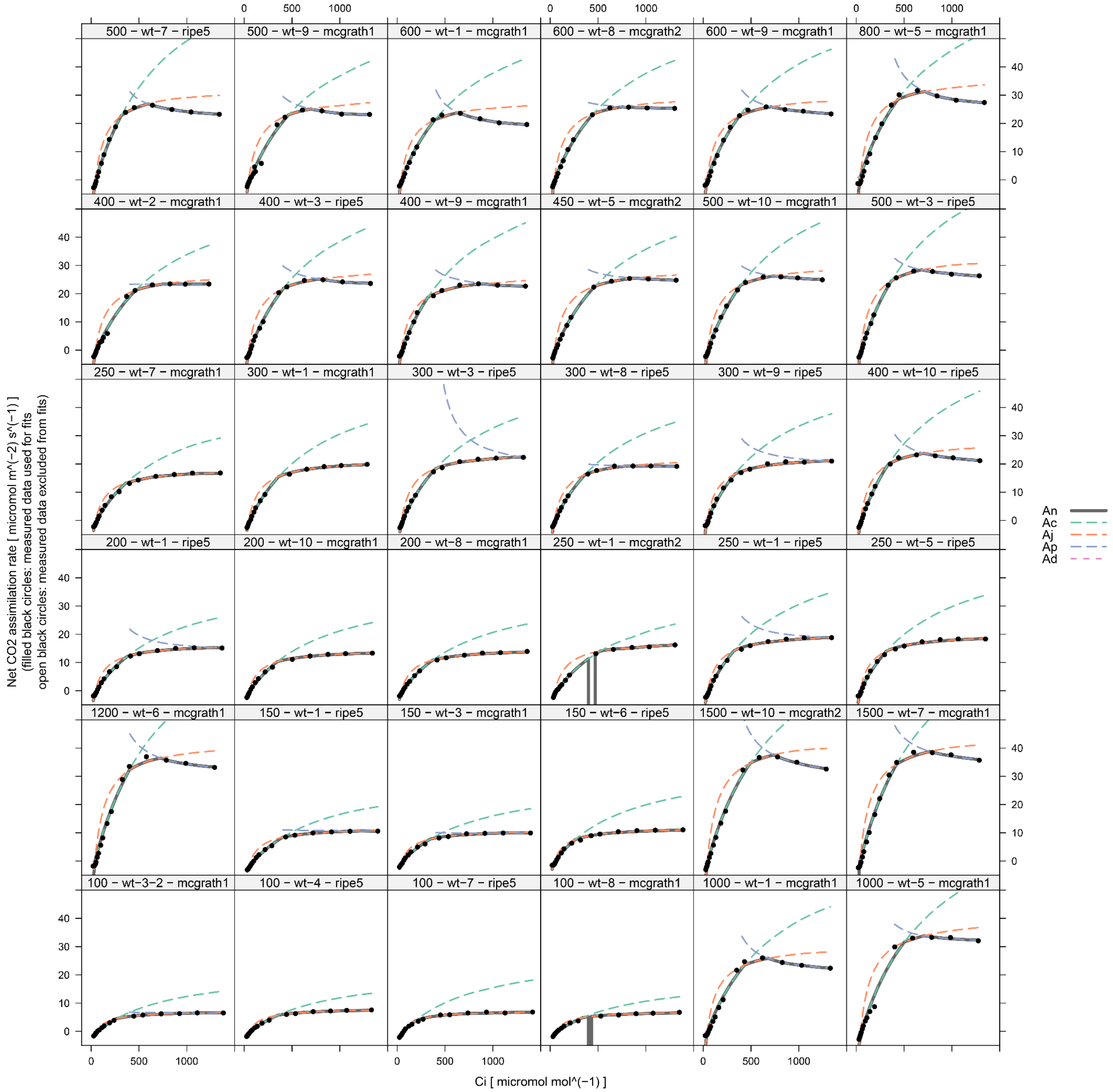

*Figure S4: Min-A+FTT variant (Equation **2**) curve fits made using all measured points, shown across the full range of $A_n$ and $C_i$.*

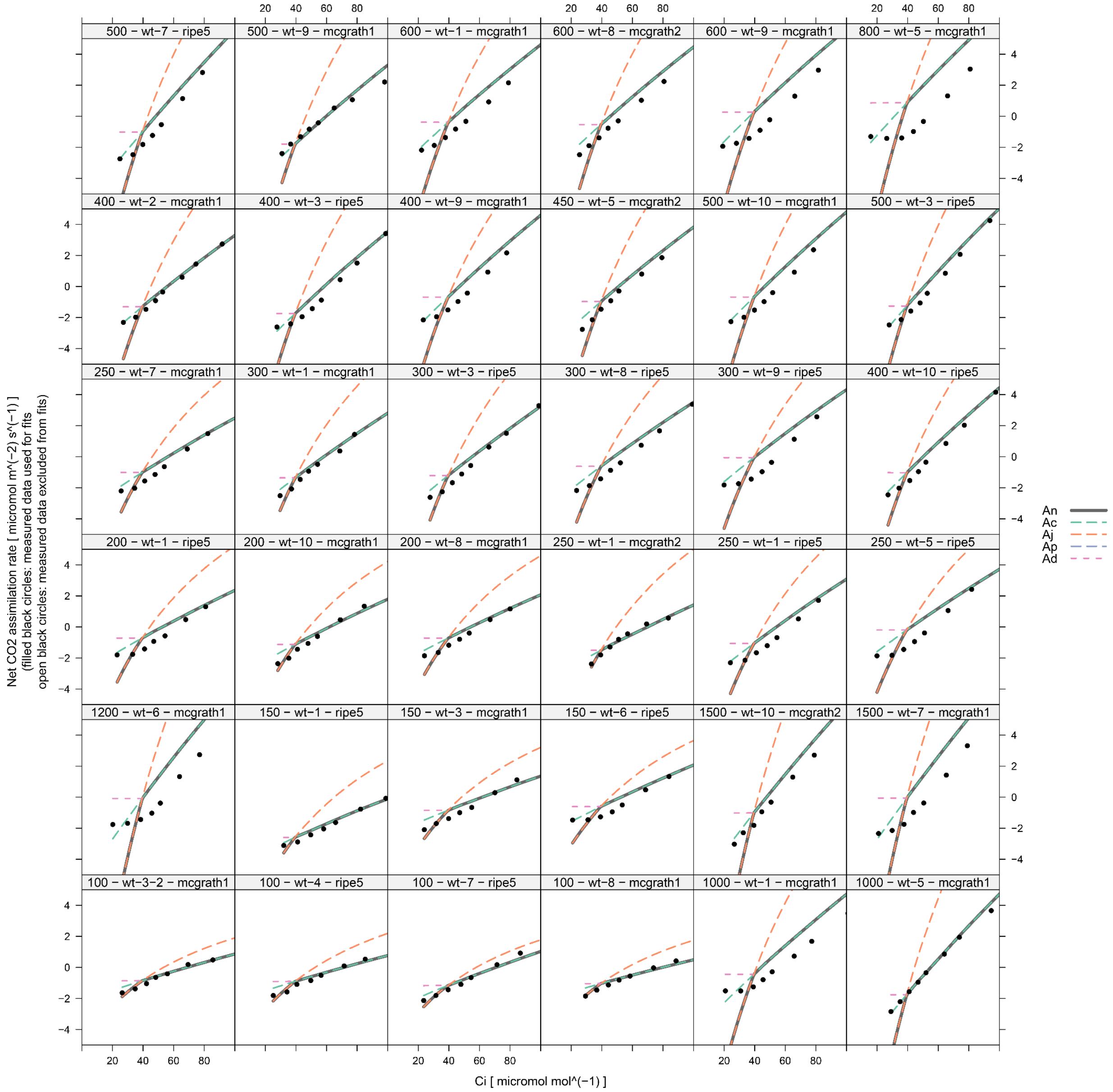

*Figure S5: Min-A+FTT variant (Equation **2**) curve fits made using all measured points, shown across a limited range of $A_n$ and $C_i$ in order to highlight the fits at low $C_i$.*

#### S5.2.1 Min-*W* Variant ($C_i$ > 45 µbar)

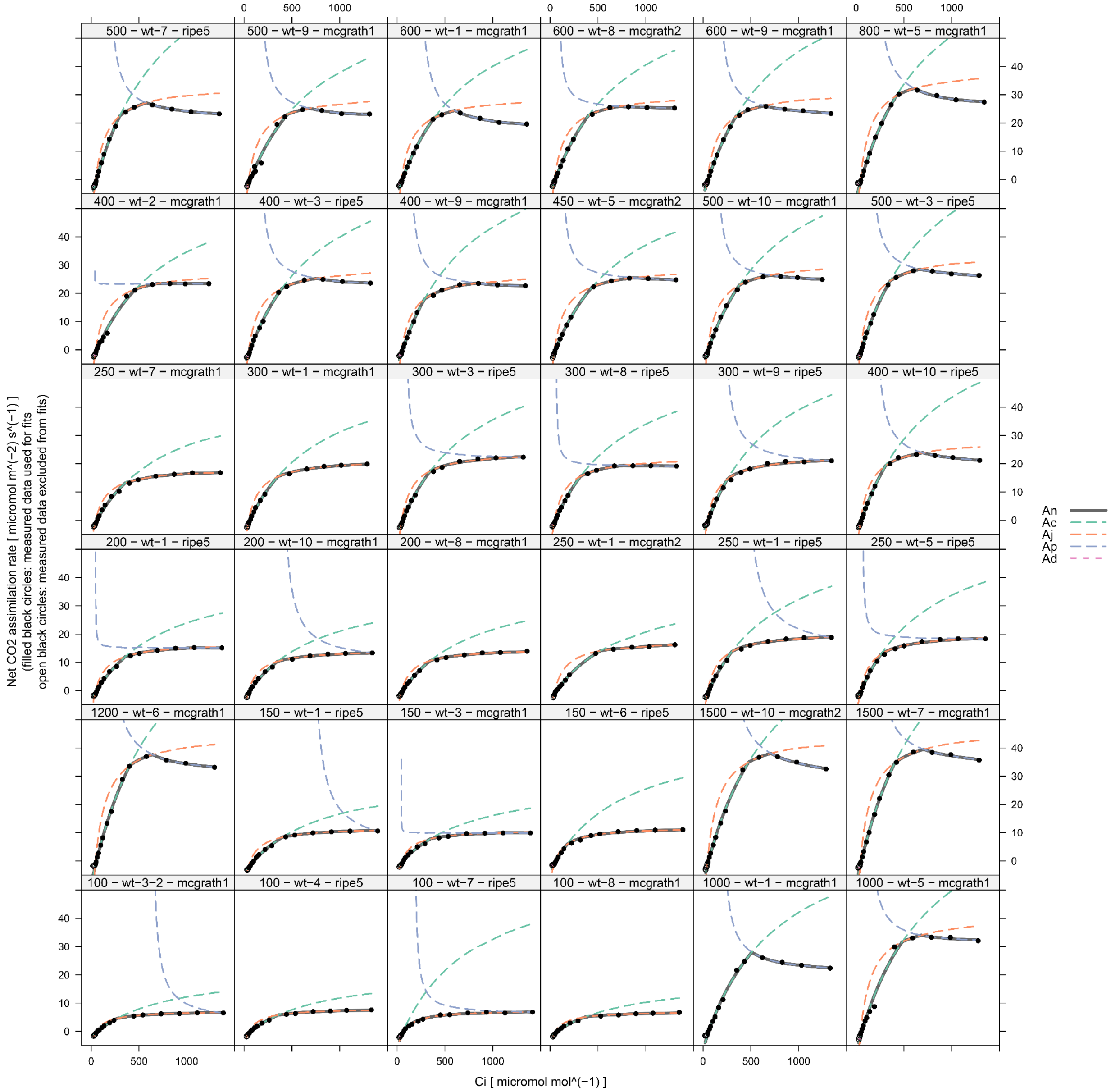

*Figure S6: Min-W variant (Equation **1**) curve fits made using measured points where $C_i$ > 45 µbar, shown across the full range of $A_n$ and $C_i$.*

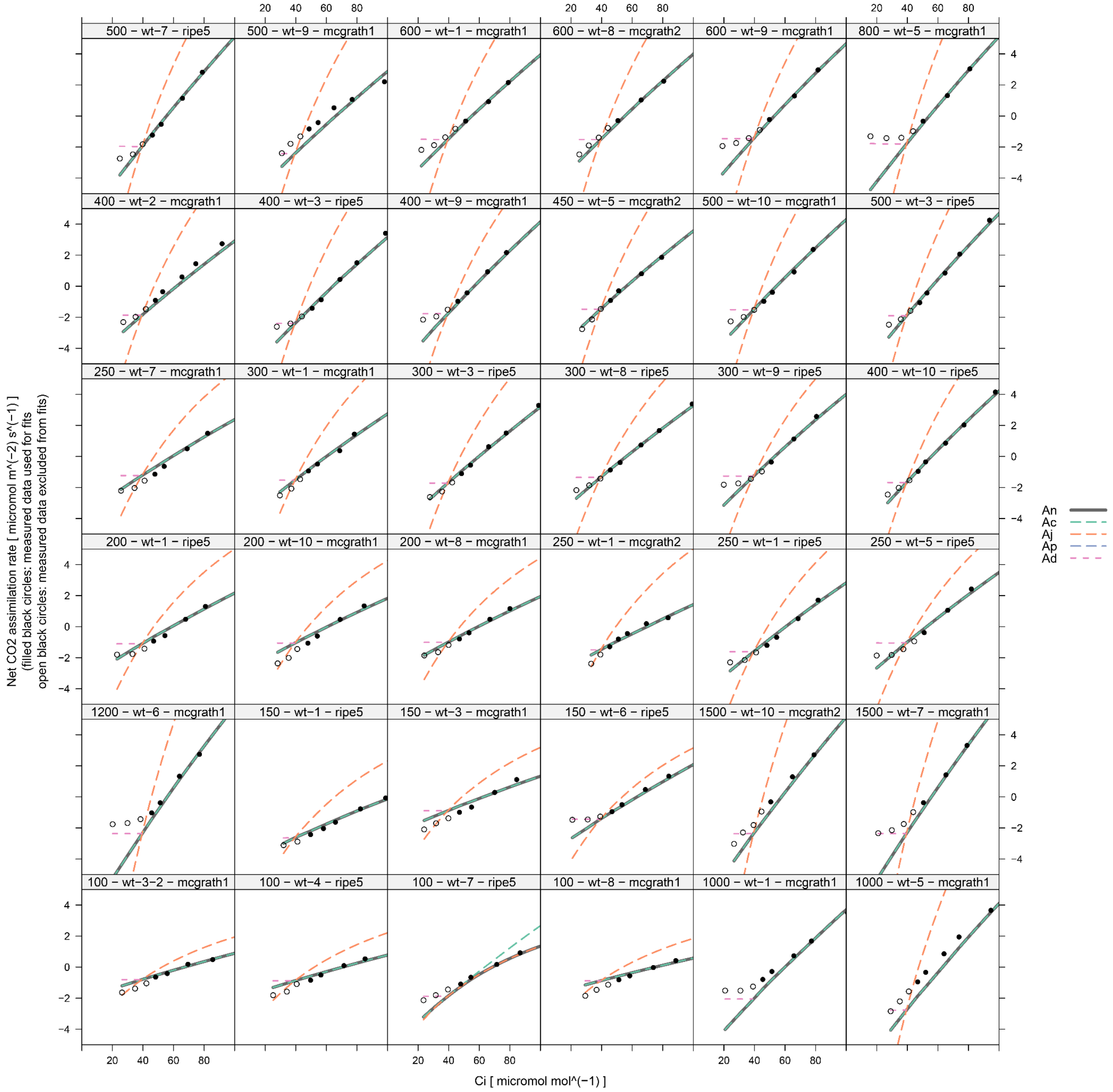

*Figure S7: Min-W variant (Equation **1**) curve fits made using measured points where $C_i$ > 45 μbar, shown across a limited range of $A_n$ and $C_i$ in order to highlight the fits at low $C_i$.*

#### S5.2.1 Min-*A*+FTT variant ($C_i$ > 45 µbar)

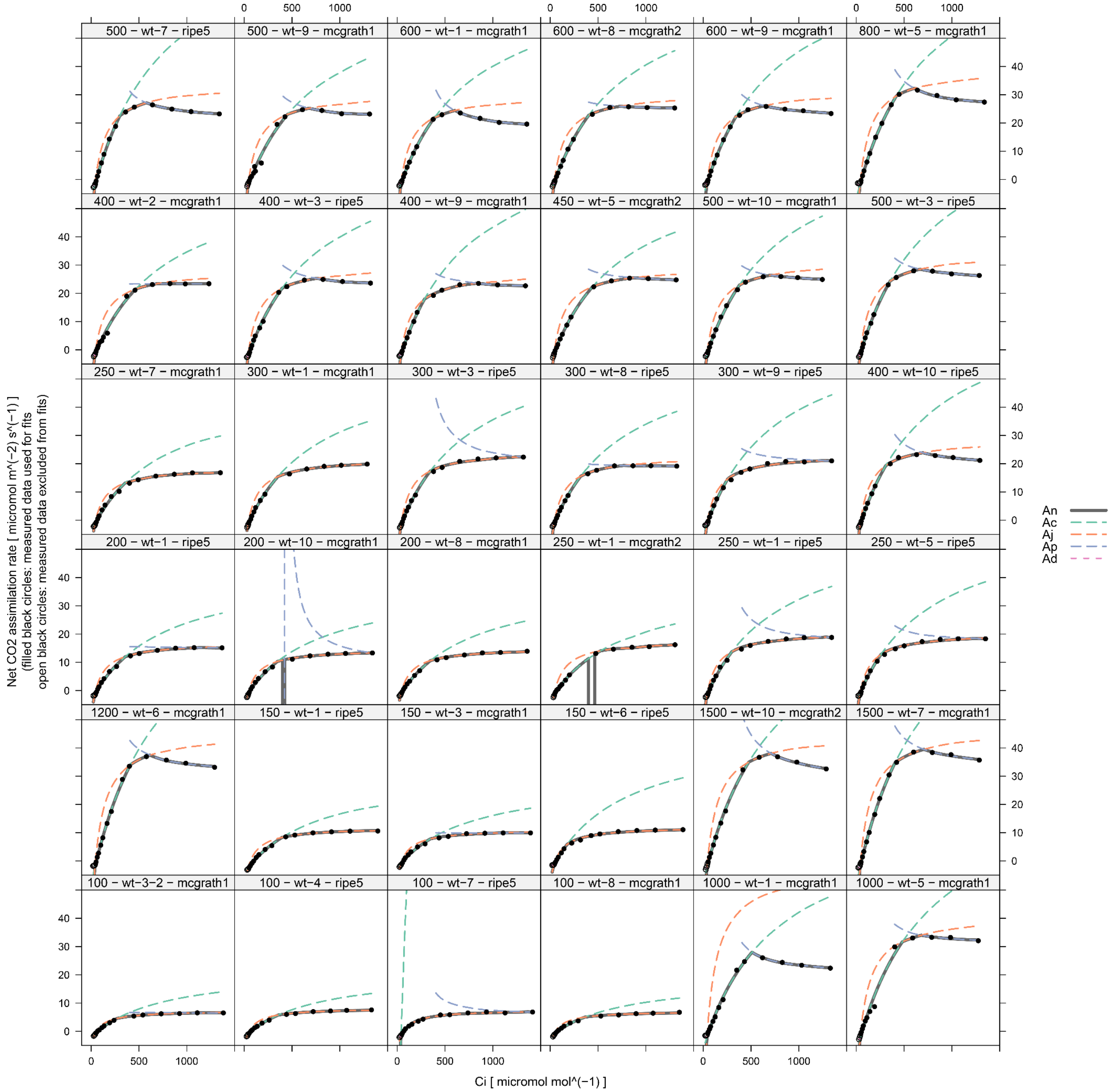

*Figure S8: Min-A+FTT variant (Equation **2**) curve fits made using measured points where $C_i$ > 45 µbar, shown across the full range of $A_n$ and $C_i$.*

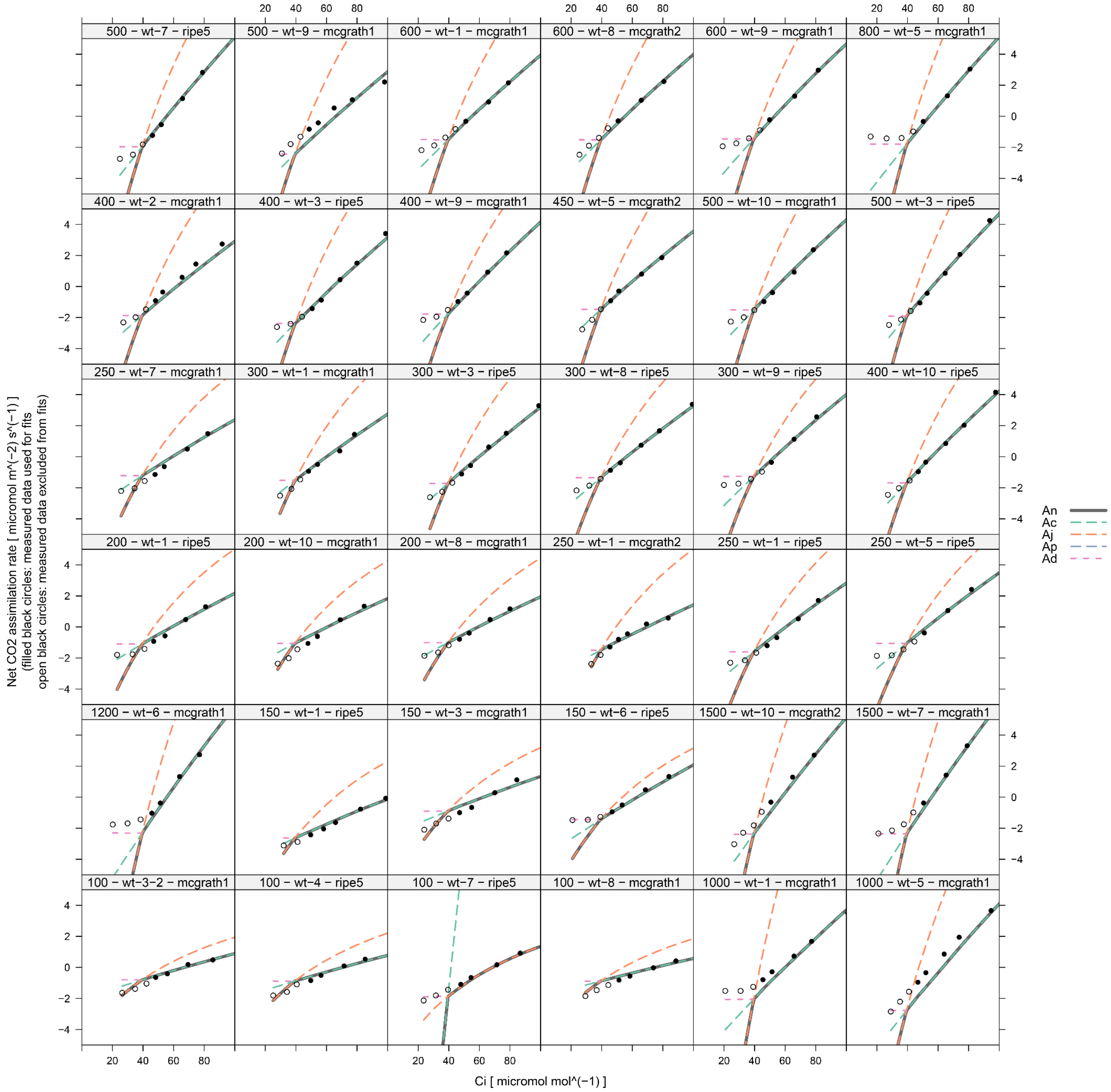

*Figure S9: Min-A+FTT variant (Equation **2**) curve fits made using measured points where $C_i$ > 45 µbar, shown across a limited range of $A_n$ and $C_i$ in order to highlight the fits at low $C_i$.*

#### S5.3 All RMSE Values

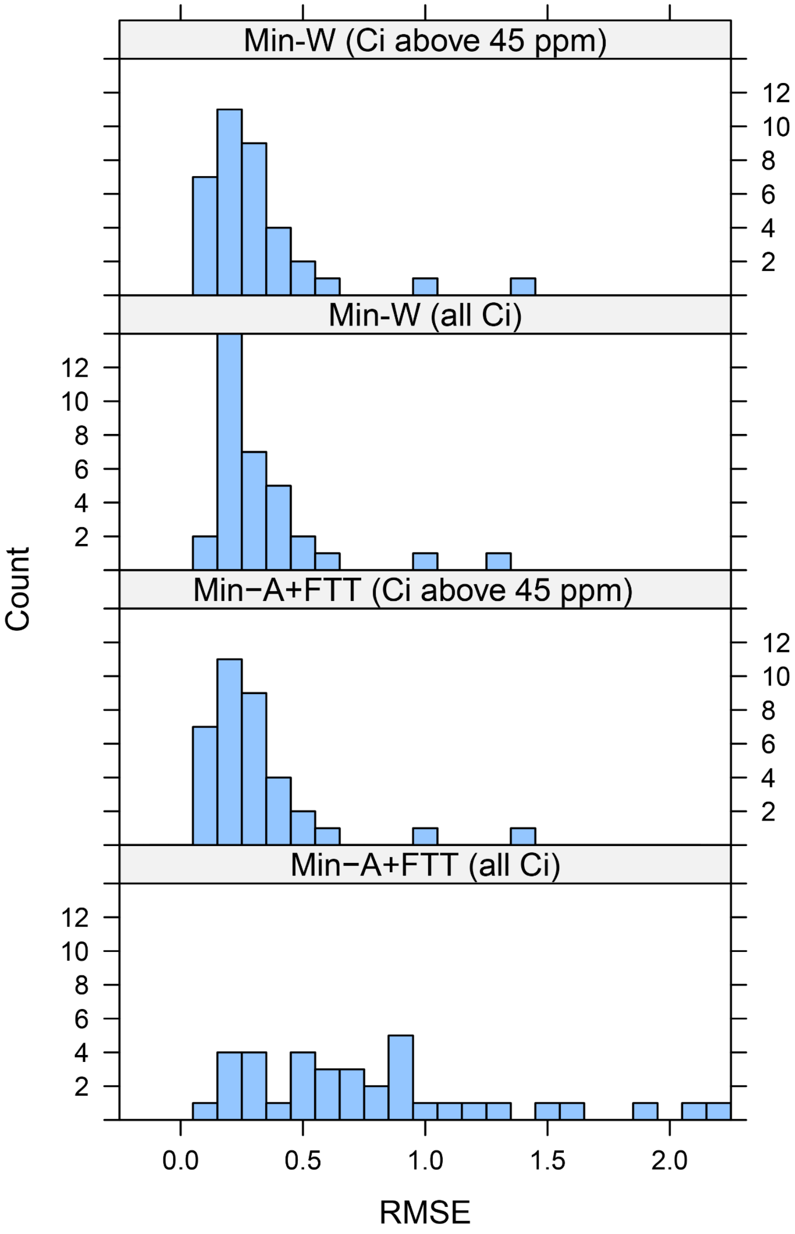

*Figure S10: Histograms of RMSE values from each fit type.*

Walker, Anthony P., Andrew P. Beckerman, Lianhong Gu, Jens Kattge, Lucas A. Cernusak, Tomas F. Domingues, Joanna C. Scales, Georg Wohlfahrt, Stan D. Wullschleger, and F. Ian Woodward. 2014. "The Relationship of Leaf Photosynthetic Traits – Vcmax and Jmax – to Leaf Nitrogen, Leaf Phosphorus, and Specific Leaf Area: A Meta-Analysis and

Modeling Study." *Ecology and Evolution* 4 (16): 3218–35. https://doi.org/10.1002/ece3.1173.

Walker, Anthony P., Abbey L. Johnson, Alistair Rogers, Jeremiah Anderson, Robert A. Bridges, Rosie A. Fisher, Dan Lu, Daniel M. Ricciuto, Shawn P. Serbin, and Ming Ye. 2021. "Multi-Hypothesis Comparison of Farquhar and Collatz Photosynthesis Models Reveals the Unexpected Influence of Empirical Assumptions at Leaf and Global Scales." *Global Change Biology* 27 (4): 804–22. https://doi.org/10.1111/gcb.15366.

Wullschleger, S. D. 1993. "Biochemical Limitations to Carbon Assimilation in C3 Plants—A Retrospective Analysis of the A/Ci Curves from 109 Species." *Journal of Experimental Botany* 44 (5): 907–20. https://doi.org/10.1093/jxb/44.5.907.

Xiao, Yi, Jen Sloan, Chris Hepworth, Colin P. Osborne, Andrew J. Fleming, Xingyuan Chen, and Xin-Guang Zhu. 2021. "Estimating Uncertainty: A Bayesian Approach to Modelling Photosynthesis in C3 Leaves." *Plant, Cell & Environment* 44 (5): 1436–50. https://doi.org/10.1111/pce.13995.

Xu, Yuan, Xinyu Fu, Thomas D Sharkey, Yair Shachar-Hill, and and Berkley J Walker. 2021. "The Metabolic Origins of Non-Photorespiratory CO2 Release during Photosynthesis: A Metabolic Flux Analysis." *Plant Physiology* 186 (1): 297–314. https://doi.org/10.1093/plphys/kiab076.

Yamori, Wataru, Kouki Hikosaka, and Danielle A. Way. 2014. "Temperature Response of Photosynthesis in C3, C4, and CAM Plants: Temperature Acclimation and Temperature Adaptation." *Photosynthesis Research* 119 (1): 101–17. https://doi.org/10.1007/s11120-013-9874-6.
